# Leucine-Rich Repeat Kinase 2 limits dopamine D1 receptor signaling in striatum and biases against heavy persistent alcohol drinking

**DOI:** 10.1101/2022.05.26.493614

**Authors:** Daniel da Silva e Silva, Aya Matsui, Erin M. Murray, Adamantios Mamais, Marlisa Shaw, Dorit Ron, Mark R. Cookson, Veronica A. Alvarez

## Abstract

The transition from hedonic alcohol drinking to problematic drinking is a hallmark of alcohol use disorder that occurs only in a subset of drinkers. This transition is known to require long-lasting changes in the synaptic drive and the activity of striatal neurons expressing dopamine D1 receptor (D1R). The molecular mechanisms that generate vulnerability in some individuals to undergo the transition are less understood. Here, we report that the Parkinson’s-related protein leucine-rich repeat kinase 2 (LRRK2) modulates striatal D1R function to affect the behavioral response to alcohol and the likelihood that mice transition to heavy, persistent alcohol drinking. Deletion of the *Lrrk2* gene specifically from D1R-expressing neurons potentiates D1R signaling at the cellular and synaptic level, enhancing alcohol-related behaviors and drinking. Mice with cell-specific deletion of *Lrrk2* are more prone to heavy alcohol drinking and consumption is insensitive to punishment. These findings identify a novel role for LRRK2 function in the striatum in promoting resilience against heavy and persistent alcohol drinking.

## Introduction

Alcohol use disorder (AUD) is a chronic relapsing disorder characterized by an inability to stop or control alcohol use despite adverse consequences (*1*). Loss of control over alcohol drinking leads to abuse and hinders long-term abstinence, driving relapse. Epidemiological data indicates there is significant individual variability in the response to alcohol and the likelihood of developing AUD (*2-4*). For example, while alcohol consumption is highly prevalent among the adult U.S. population, only a fraction (∼ 8%) of those who consume alcohol are diagnosed with AUD each year (*5*). These and other data point to the existence of risk and resilience factors for developing the disorder.

The leucine-rich repeat kinase 2 (*Lrrk2)* gene is highly expressed in the striatum where it plays an important role in regulating synapse formation and synaptic transmission (*6*). Mutations in the human *LRRK2* gene are associated with Parkinson’s disease (*7, 8*), which is characterized by prominent impairment in dopamine signaling and basal ganglia function. Recent work has linked changes in striatal expression of the *Lrrk2* gene with alcohol drinking in humans and rodents (*9, 10*). We recently reported a positive correlation between striatal levels of *Lrrk2* mRNA and alcohol drinking levels in mice, especially an association with inflexible alcohol drinking that persists despite adverse outcomes or punishment (*9*).

The striatum plays a central role in the learning and execution of reward-motivated and goal-directed behaviors (*11*). It is also involved in the process of action selection (*12*), which highlights the relevance of the basal ganglia in neuropsychiatric disorders such as substance use disorders. Functional and morphological alterations in the striatal circuitry have been linked to not only AUD but also attention deficit hyperactivity disorder, obsessive-compulsive disorder, and substance use disorders (*13-15*). At the cellular level, the striatum is composed of two main classes of projection neurons: D2R-expressing medium spiny neurons (D2-MSN) and D1R-expressing medium spiny neurons (D1-MSN). These neurons express distinct dopamine receptors, project to different outputs in the basal ganglia, and regulate opposing processes and behaviors. D1-MSNs are known to form the striatonigral direct-pathway or go-pathway because of their role in reinforcement and reward learning, while D2-MSNs are referred to as striatopallidal indirect pathway or no-go pathway because of their role in avoidance and aversive learning (*16, 17*).

Recent data indicate that LRRK2 modulates D1R function via regulation of receptor internalization and the subcellular localization of Protein kinase A (PKA), the main signaling molecule downstream of D1R activation (*18-20*). For example, mutations in the *Lrrk2* gene alter membrane trafficking and surface expression of D1R (*20, 21*). Global deletion of *Lrrk2* in mice enhanced PKA signaling downstream of D1R and altered dendritic spine morphology and synaptic strength in striatal D1-MSNs (*18, 19*). Gain-of-function mutations of *Lrrk2* were shown to reduce PKA activity (*18*). Thus, these published studies suggest that the LRRK2 acts as a negative modulator of D1R signaling in the striatum.

D1R and D1-MSNs in the striatum regulate the reinforcing properties of alcohol. Further, alcohol exposure elicits functional and structural plasticity selectively in striatal D1-MSNs. For example, a single alcohol drinking session potentiates synaptic drive onto D1-MSNs in the nucleus accumbens (*22*). Longer exposure to alcohol also triggers enduring changes in the circuitry of the dorsomedial striatum (DMS). Repeated cycles of alcohol exposure induce a long-lasting potentiation in excitatory synaptic transmission in D1-MSNs in the DMS, but not in D2-MSNs (*23*). Additionally, the deletion of the D1R gene (*Drd1a*) impairs alcohol drinking and preference (*24*). Pharmacological blockade of D1-like receptors in the DMS, but not D2R, attenuates alcohol consumption (*22, 23, 25*). Further, D1-like receptor antagonist impair alcohol-seeking in an operant task (*26*). Collectively, these findings point to an important role of D1R function and striatal D1-MSN activity in driving alcohol drinking. We then hypothesize that enhanced D1R function and recruitment of striatal D1-MSN activity during alcohol exposure is a possible mechanism contributing to AUD vulnerability.

Here, we test the hypothesis that LRRK2 functions to lower D1R function in the direct-pathway D1-MSNs and therefore limits alcohol reinforcing properties. Under this hypothesis, LRRK2 activity in D1-MSNs confers individuals with resilience against heavy alcohol drinking and drinking despite adverse consequences. Using mice carrying conditional *Lrrk2* alleles, we generated cell-specific deletion of the *Lrrk2* gene in D1-MSNs or D2-MSNs and evaluated D1R function and alcohol-related behaviors. The results show that selective deletion of *Lrrk2* in D1-MSNs, but not D2-MSNs, potentiates the cellular and behavioral response to D1-like receptor agonist and alcohol. Loss of *Lrrk2* in D1-MSNs promotes heavy and punishment-resistant alcohol drinking, suggesting that LRRK2 activity in direct-pathway neurons is an important factor that determines vulnerability to out-of-control alcohol drinking.

## Results

### Lrrk2 expression is enriched in Medium Spiny Neurons in the striatum

We evaluated the expression profile of *Lrrk2* mRNA in different brain regions associated with reward learning and reinforcement using public available RNA-seq dataset (*27-29*). We found that *Lrrk2* mRNA is remarkably enriched in the striatum (ventral and dorsal) compared to other brain regions, such as the hippocampus, prefrontal cortex, basolateral amygdala, and ventral tegmental area (Fig. 1A). Using RNA *in situ* hybridization we investigated the cellular specificity of *Lrrk2* mRNA expression in D1-MSNs and D2-MSNs, the two most common neuronal types in the striatum (Fig. 1B). Similar median values of *Lrrk2* mRNA puncta were measured in cells that were co-labeled with *Drd1a* and with *Adora2a*, markers for D1-MSNs and D2-MSNs, respectively (Fig. 1C; Mann-Whitney U=42802, *p*=0.74). Approximately 95% of identified D1-MSNs (Drd1a+) and 92% of identified D2-MSNs (Adora2a+) showed at least one *Lrrk2* mRNA puncta (Fig. 1D).

**Figure 1:**
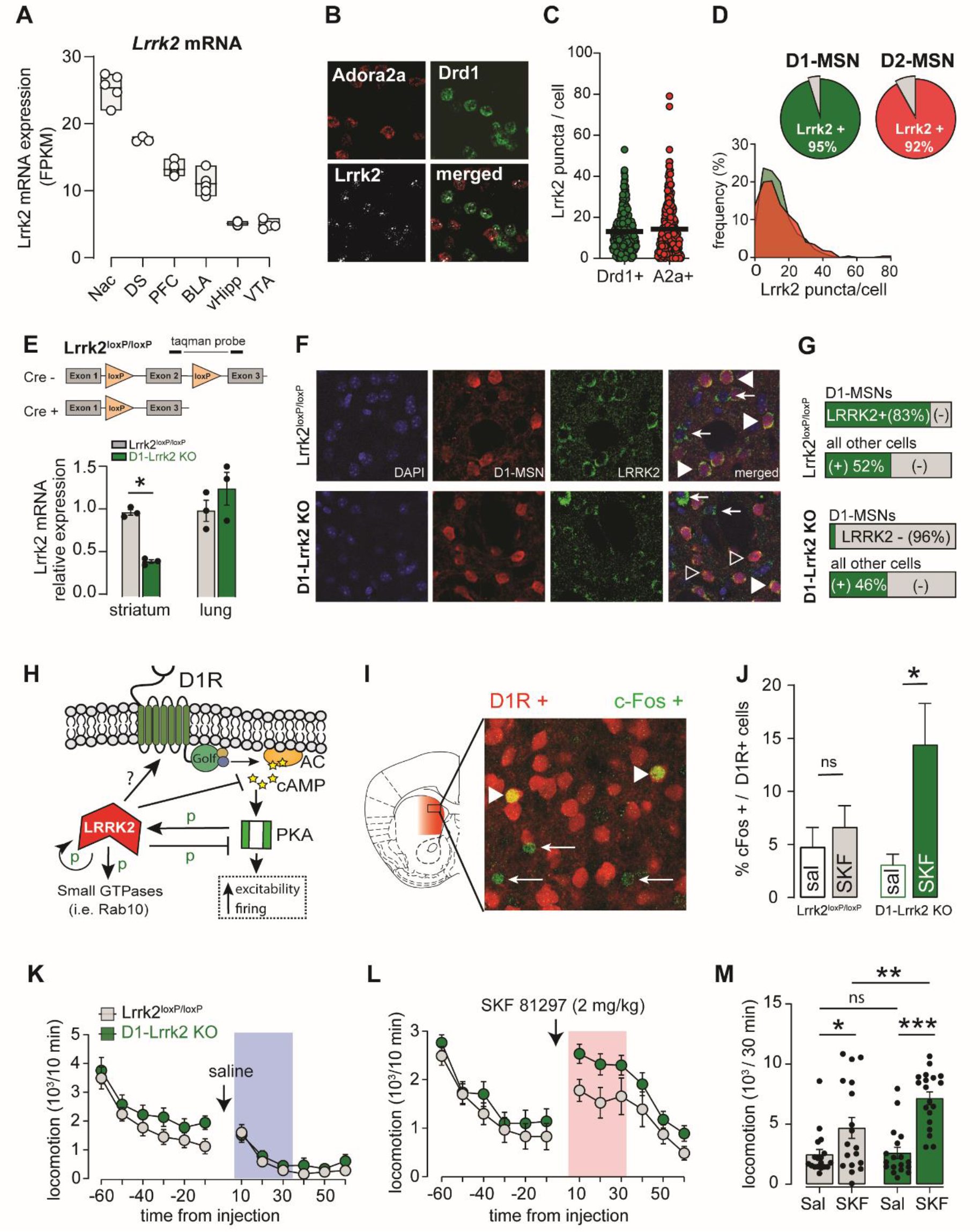
Targeted deletion of *Lrrk2* to D1R-expressing neurons potentiates the behavioral and cellular response to dopamine D1R activation. **A**, *Lrrk2* mRNA expression across brain nuclei of the mesolimbic cortical circuit. Abbreviations: NAc, nucleus accumbens; DS, dorsal striatum; PFC, prefrontal cortex; BLA, basolateral amygdala; vHipp, ventral hippocampus; VTA, ventral tegmental area. **B**, Confocal images of the dorsal striatum showing fluorescent RNA in situ hybridization with probes for *Lrrk2* (white puncta), Adora2a (red puncta), and Drd1a mRNA (green puncta). **C**, Quantification of *Lrrk2* mRNA puncta per cell that co-labels with Drd1 mRNA (green, n=271) and with Adora2a mRNA (red, n=321). Symbols represent individual values, and the line shows the median of each group. **D**, *Top*, pie chart showing the percentage of D1R-positive and A2a-positive cells that express *Lrrk2* mRNA in the dorsal striatum. *Bottom*, frequency distribution of *Lrrk2* mRNA puncta in D1R-positive and A2a-positive cells in the dorsal striatum. **E**, *Top*, Diagram of the conditional *Lrrk2* allele showing the LoxP insertion site. *Bottom*, Real-time qPCR of *Lrrk2* mRNA expression in striatum and lung. **F**, Confocal images of fluorescence immunostaining for endogenous LRRK2 protein (green), tdTomato in D1-MSNs (red), and DAPI (blue) in DMS sections from D1-Lrrk2-KO mice and controls. Filled arrowheads point to cells co-labeled with LRRK2 and tdTomato; open arrowheads point to cells labeled with tdTomato and negative for LRRK2; arrows point to cells labeled with LRRK2 and negative for tdTomato. **G**, Quantification of the degree of LRRK2 protein co-labeled with tdTomato (D1-MSNs) or without it (D1-negative). **H**, Modulation of D1R signaling and function by LRRK2 kinase activity. G_olf_, guanine nucleotide-binding protein G_(olf)_ subunit alpha; cAMP, cyclic adenosine monophosphate; AC, adenylate cyclase; PKA, protein kinase A. **I**, Representative confocal image of DMS from D1-Lrrk2-KO mouse showing red fluorescence in D1R positive cells and green fluorescence and c-Fos immunolabeling following injection of SKF81297. Arrows point to c-Fos positive cells and arrowheads point to double positive cells (c-Fos+;tdTomato). **J**, Quantification of double positive cells (c-Fos+;tdTomato) as percent of total D1R-expressing cells (tdTomato+) after systemic administration of saline or SKF81297 (2 mg/kg). **K**, Basal locomotion measured before and after saline injection. **L**, Locomotion measured before and after SKF81297. **M**, Average locomotor activity during the initial 30 minutes post saline and SKF81297. For all panels, bars represent mean ± S.E.M and symbols represent values from individual mice. (*) denotes *P* < 0.05, ** P < 0.01, *** *P* < 0.001.

### Deletion of *Lrrk2* from direct-pathway D1-MSNs potentiates D1R-like function

Because LRRK2 acts as a negative modulator of D1R function in the striatum (*18, 19*), we hypothesized that loss of LRRK2 activity in D1-MSN potentiates D1R function and promotes behavioral output associated with the direct-pathway. We crossed mice bearing conditional alleles for *Lrrk2* gene (Lrrk2^loxP/loxP^) with mice expressing Cre recombinase under the *Drd1a* promoter to generate a transgenic mouse line with targeted deletion of the *Lrrk2* gene to D1R-expressing neurons (D1-Lrrk2-KO, Fig. 1E). qPCR analysis showed 60% reduction of *Lrrk2* mRNA levels in striatal samples from D1-Lrrk2-KO mice compared to their littermate Lrrk2^loxP/loxP^ controls (Fig 1E; t_(4)_=14.5 p<0.001) and no differences the lung, a tissue that expresses *Lrrk2* but not *Drd1a* (Fig 1E; t_(4)_=1.13, p=0.32). Immunocytochemistry was used to validate the cell-specificity of LRRK2 deletion at the protein level. D1-Lrrk2-KO mice were crossed with the Drd1-tdTomato line to fluorescently label D1-MSNs. Staining for LRRK2 was observed in 83% of D1R-positive cells in the DMS of control mice whereas only in 4% of D1R-positive cells in D1-Lrrk2-KO mice (Fig. 1G; *X*^*2*^_(1,398)_=249, *P*<0.0001). Importantly, the percentage of D1R-negative cells that expressed LRRK2 was similar in both genotypes (Fig. 1G; 52% *vs* 46%; *X*^*2*^_(1,634)_=2.2, *P*=0.14).

LRRK2 is thought to regulate D1R function via modulation of intracellular signaling (Fig. 1H). We tested whether *Lrrk2* deletion in D1-MSNs alters D1R induction of the immediate-early gene c-Fos in the striatum (Fig. 1I). A low dose of the D1-like receptor agonist SKF81297 (2 mg/kg, *i*.*p*.) or saline was administered to D1-Lrrk2-KO and littermate Lrrk2^flox/flox^, both crossed into a Drd1-tdTomato background. There was no general difference in the percentage of D1-MSNs labeled with c-Fos in the DMS of control and D1-Lrrk2-KO mice (Fig. 1J; no genotype effect: F_(1,59)_=1.7, *P*=0.2), however, low dose of SKF81297 produced a significant increase in the percent of c-Fos expressing D1-MSNs specifically in D1-Lrrk2-KO mice and no changes in controls when compared to saline treated mice (Fig. 1J; SKF effect: F_(1,59)_=8.2, p<0.01; interaction: F_(1,59)_=4.2, *P*<0.05; Sidak’s test *P*<0.01 for D1-Lrrk2-KO and *P*=0.8 for controls). We assessed whether postnatal deletion of *Lrrk2* was sufficient to enhance D1R-mediated c-Fos induction by expressing Cre(eGFP) recombinase or GFP in the DMS of adult Lrrk2^flox/flox^ mice. Five weeks post-surgery, mice were injected with either saline or SKF81297 (2 mg/kg, *i*.*p*.) and brains were processed for c-Fos immunostaining. There was no general group differences in the number of c-Fos positive cells in the DMS of Cre(eGFP)-injected and GFP-injected mice (Fig S1A; no Cre effect: F_(1,40)_=0.72, *P*=0.4). SKF81297 injection did not increase the percentage of c-Fos labeled cells in neither Cre(eGFP)-injected nor GFP-injected mice. (Fig S1A; no SKF effect: F_(1,40)_=0.02, *P*=0.9). Thus, post-natal deletion of the *Lrrk2* gene, within 5 weeks, does no produce an enhancement in D1R response and therefore early developmental deletion of *Lrrk2* is required for the D1R hypersensitivity.

The behavioral impact of *Lrrk2* deletion, was assessed by recording locomotor activity after injection of saline or SKF81297 (2 mg/kg, *i*.*p*.) in naïve mice. Baseline locomotion measured before and after saline injection was similar between genotypes (Fig. 1K; pre-injection: F_(1,35)_=2.3, *P=*0.13; post-injection: F_(1,35)_=0.79, *P*=0.38). SFK81297 administration increased locomotion in both genotypes (Fig. 1L-M; REML, SKF: F_(1,34)_=34.8; genotype: F_(1,3)_=4.3; interaction: F_(1,34)_=4.2, *P*<0.05), however, D1-Lrrk2-KO mice showed larger locomotor stimulation during the first 30 min after injection, indicating an enhanced behavioral response to D1-like agonist (Sidak’s test *P*<0.01 for SKF control vs SKF D1-Lrrk2-KO). Thus, these findings provide evidence of enhanced cellular and behavioral response to D1R following the constitutive loss of LRRK2 function in D1-MSNs.

### Enhanced physiological response to D1-like agonist in striatal D1-MSNs

Under specific conditions, D1R activation can increase the excitability of striatal D1-MSNs (*30-32*) and increase GABA transmission from D1-MSN terminals in the substantia nigra pars reticulata (SNr)(*33, 34*). Thus, we assessed physiological readouts of D1R activation using whole-cell recordings from D1-MSNs in the DMS. In current-clamp recordings, input-output curves under baseline conditions showed a similar number of action potentials fired as a function of current injected in D1-MSNs of both genotypes (Fig. 2A, B; no genotype effect: F_(1,31)_=0; *P*=0.96; n=15-18 cells, 7/7 mice). However, after incubation with SKF81297 (1μM), there was a leftward shift in the input-output curve from D1-Lrrk2-KO (Fig. 2A,C; genotype effect: F_(1,41)_=4.1; *P*<0.05; interaction: F_(6,246)_=4.7, *P*<0.0001; n=21-22 cells, 7/7 mice). A cumulative histogram of latency to the first spike at 500pA current step showed a shift to shorter latencies after SKF81297 application in D1-Lrrk2-KO (Fig. 2E; t_(38)_=2, p=0.052) and no change in control mice (Fig. 2D; t_(33)_=0.1, p=0.91). We also found a strong trend towards an increase in the number of spikes within the first half of the current step and in D1-Lrrk2-KO mice (Fig. 2F; genotype effect: F_(1,73)_=3.8, *P*=0.054). These results suggest that D1-like agonist shortened the latency to spike and increased firing of action potentials right after depolarization in D1-MSNs lacking LRRK2. Electrically evoked AMPAR/NMDAR ratio was not different between genotypes (Fig. S1B; t_(18)_=0.01, *P*=0.99, n=10/10 cells, 4/3 mice), suggesting no significant changes to the excitatory drive to D1-MSNs.

**Figure 2:**
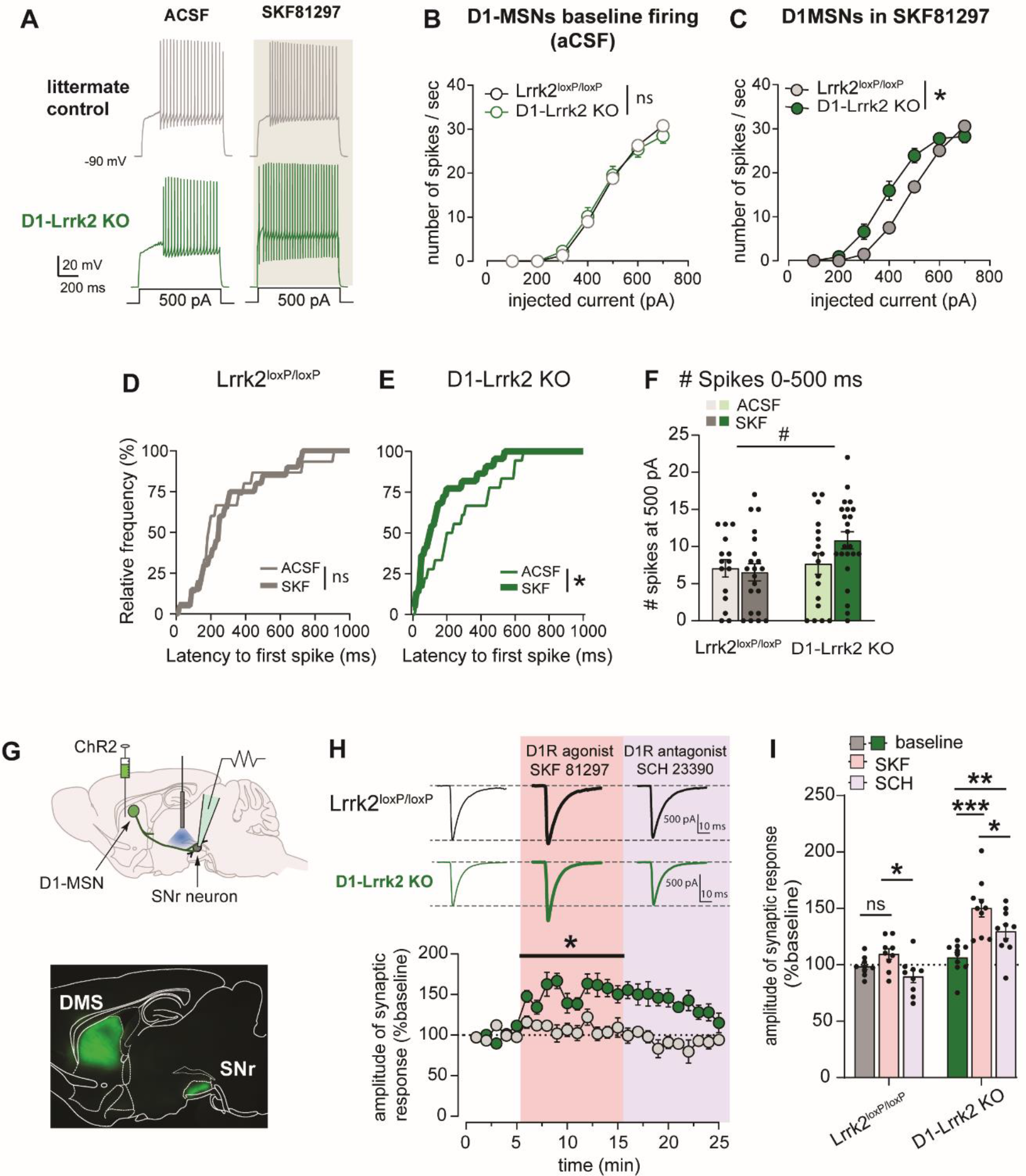
Deletion of *Lrrk2* in D1R-expressing neurons potentiates the electrophysiological response to D1-like agonist in these cells. **A**, Representative traces of action potentials recorded from D1-MSNs in response to current step injection during whole-cell current-clamp recordings in control aCSF conditions and after incubation with SKF81297 (1 μM) in the DMS of D1-Lrrk2-KO (green) mice and littermate controls (gray). **B-C**, Input-output curve of firing rate of D1-MSNs in response to current steps of increasing amplitude after incubation of slices with aCSF (B) or SKF81297 (C). **D-E**, Cumulative histogram of the latency to the first action potential at 500pA current step for littermate controls (D) and D1-Lrrk2-KO mice (E). **F**, Number of action potentials during the first half (0-500ms) of the 500pA current step. **G**, *Top*, schematic diagram showing injection site of ChR2 viral vector in the DMS, the site of the recordings in the midbrain substantia nigra reticulata (SNr), and the local fiber optic used to stimulate direct-pathway axons in the SNr; *Bottom*, fluorescent image shows expression of ChR2 (green) in the DMS and labeled axons in the SNr. **H**, *Top*, representative traces of voltage-clamp recordings from neurons in the SNr showing optogenetic-evoked synaptic response during baseline and after bath application of SKF81297 and SCH23390 in brain slices from D1-Lrrk2-KO (green) and littermate controls (gray); *Bottom*, time course of recorded inhibitory synaptic responses in putative GABAergic neurons of SNr in response to optogenetic stimulation of direct-pathway MSNs in the presence of SKF81297 (pink shaded area) or SCH23390 (purple shaded area). **I**, Average of the normalized oIPSC amplitude (first 5 minutes) in response to SKF81297 and SCH23390 application. For all panels, bars represent mean ± S.E.M and symbols represent values from individual slices. (*) denotes *P* < 0.05, (***) denotes *P* < 0.001.

D1-MSNs in the dorsal striatum send axonal projections to the substantia nigra reticulata (SNr) (*35, 36*). D1Rs are localized to the synaptic terminals of these neurons where they are thought to potentiate GABA transmission. We expressed Channelrhodopsin-2 in the DMS and used optogenetic stimulation to evoke GABA transmission while recording from SNr neurons (Fig. 2G). Whole-cell recordings showed reliable optical-evoked inhibitory postsynaptic currents (oIPSC) which were blocked by GABA-A receptor blockers (not shown). Bath application of D1-like agonist SKF81297 (1 μM) produced a 10 ± 5 % increase of oIPSC amplitude in slices from control animals, which was not statistically significant. In contrast, the same concentration of SKF81297 produced a significant 50 ± 8 % increase in oIPSC amplitude D1-Lrrk2-KO mice (Fig. 2H,I; genotype: F_(1,17)_=34; treatment: F_(2,34)_=34; interaction: F_(2,34)_=6.4, *P*<0.005; Sidak’s test: baseline *vs* SKF: P<0.0001 for D1-Lrrk2-KO). The increase was partially reversed by subsequent application of the D1-like antagonist SCH23390, indicating the requirement for D1R activation and suggesting possible long-term actions (Fig. 2H,I; Sidak’s test: baseline vs SCH, *P*<0.05). Furthermore, SKF81297 reduced the paired-pulse ratio (PPR) in D1-Lrrk2-KO mice from 1.5 ± 0.1 to 1.2 ± 0.08 while had no effect in littermate control mice (Fig. S1C; interaction: F_(1,15)_=16, *P*<0.001; Sidak’s test: control, *P*=0.26; D1-Lrrk2-KO, *P*<0.005). Altogether, these results indicate that deletion of *Lrrk2* in D1R-expressing neurons potentiates dopamine signaling via D1 receptors facilitating GABA transmission.

### Overall unchanged dopamine-related behaviors in mice with Lrrk2 deletion in D1-MSN

Striatal D1R are important mediators of dopamine neurotransmission and are involved in the regulation of basic functions and behaviors, such as feeding, motivation, and exploration. Here, we assessed exploratory behavior, the response to novelty, food, and sweet consumption, and compared D1-Lrrk2-KO and control mice. Mice showed similar distance traveled and velocity over a 30-min period in an open-field arena (Fig 3A-B; distance: t_(14)_=0.53, *P*=0.6; velocity: t_(14)_=0.55, *P*=0.59). Time spent in the center of the arena was also similar across genotypes (Fig 3C; t_(14)_=0.6, *P*=0.56). The response to novelty was assessed as time spent exploring a novel object placed in the center of the arena over a 15-min session. D1-Lrrk2-KO and littermate control mice spent similar time exploring the novel object (Fig 3D; t_(14)_=0.8, *P*=0.42). These data suggest that general exploratory behaviors are unaffected by the deletion of the *Lrrk2* gene from D1-MSN.

**Figure 3:**
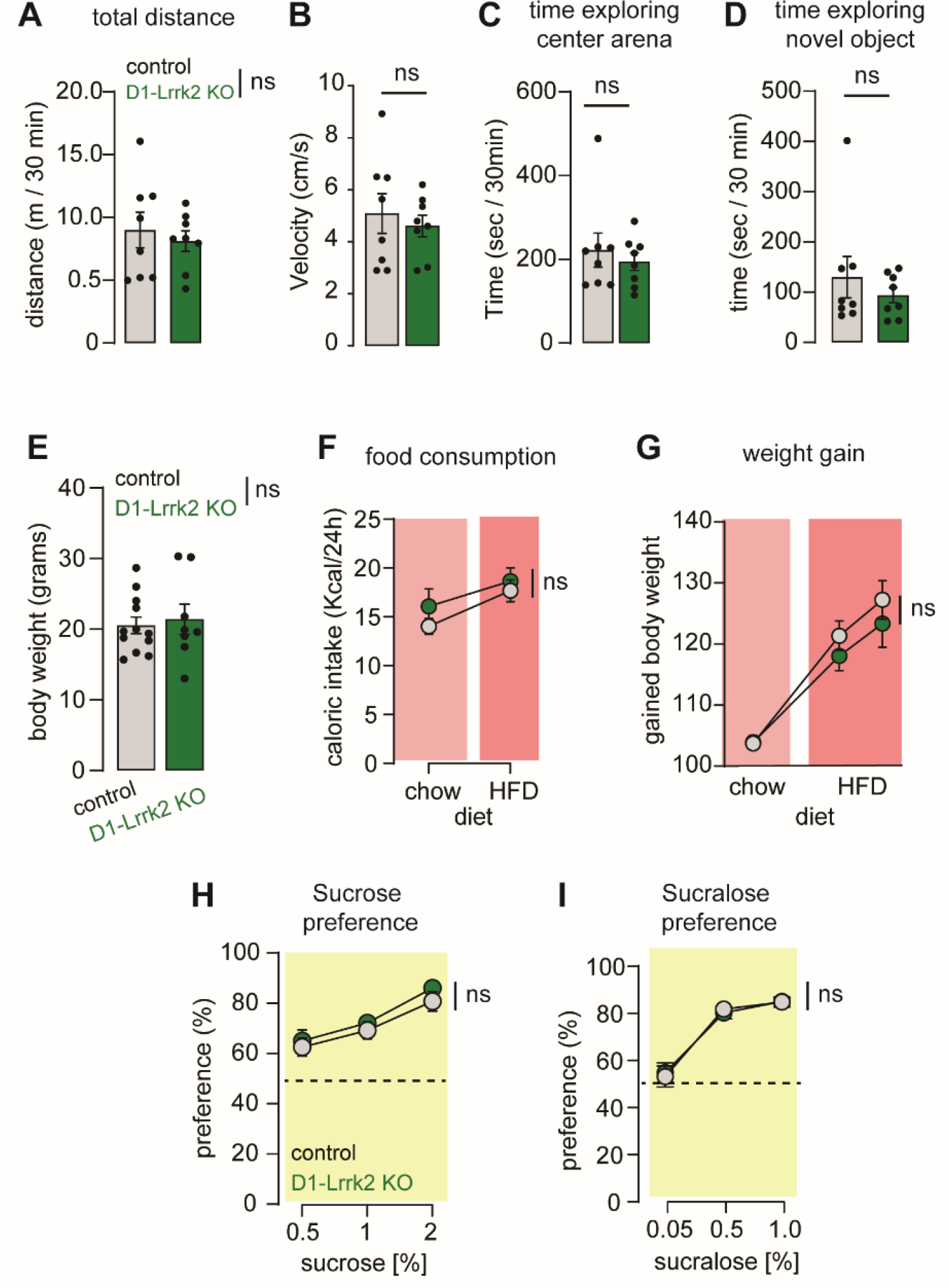
Exploration and caloric intake are largely unchanged following selective deletion of *Lrrk2* in D1-MSN. **A-D**, Distance travelled, velocity, time in center and novel object exploration assessed in an open field arena. **E**, average body weight for age-matched mice of each genotype. **F**, Caloric intake in Kcal/day when given unrestricted access to normal chow diet and high fat diet (HFD) for littermate control (gray) and D1-Lrrk2-KO mice (green). **G**, change in body weight during access to regular chow diet or HFD diet **H**. Sucrose preference during 0.5%, 1% and 2% sucrose solution access using a two-bottle choice procedure. **I**, Sucralose preference using two-bottle choice procedure.

Food consumption was measured for 2 weeks while mice had access to regular chow followed by 2 weeks when mice had access to a highly palatable high-fat diet (60 kcal% fat, HFD). At the start of the experiment, mice from both genotypes had similar body weights (Fig 3E; 21.4 ± 2.1 vs 20.1 ± 1.2 g; t_(18)_=0.4, *P*=0.7). There was no difference in the caloric intake between genotypes under either diet (Fig 3F; F_(1,18)_=1.3, *P*=0.26). There was a main effect of diet as mice from both genotypes consumed more calories under HFD compared to chow (Fig 3F; F_(1,18)_=6.8, *P*<0.05). Weight gain during different treatments was also similar for both genotypes (Fig 3G; no genotype: F_(1,18)_=0.6, *P*=0.44; no interaction: F_(2,36)_=0.7, *P=*0.7).

Preference for sucrose was assessed using a two-bottle choice procedure with 0.5, 1, and 2% sucrose solution. Overall, there was no difference in sucrose preference across genotypes when two large cohorts of mice were tested (Fig. 3H; n=22/22 mice). On the first cohort, D1-Lrrk2-KO mice showed higher preference for 1% sucrose (genotype effect: F_(1,20)_=5.7, p<0.05, Sidak’s test: 1% sucrose p<0.05; n=11/11 mice). The difference was not reproduced in a second independent cohort of mice which included more concentrations of sucrose solution (REML, no genotype effect: F_(1,42)_=1.7, *P*=0.2; dose effect: F_(2,62)_=18, *P*<0.0001, n=22/22). Last, we tested sweet taste sensitivity by measuring the preference for the non-caloric sweetener sucralose in a separated cohort of mice. Both genotypes showed a similar preference for sucralose at all concentrations (Fig 3I; concentration effect: F_(1.35,44.8)_=68, *P*<0.0001; no genotype effect: F_(1,33)_=0.001, *P*=0.97). Thus, there is no major changes in consummatory behaviors nor exploration in mice lacking the *Lrrk2* gene in D1-MSNs.

### Loss of LRRK2 function in D1-MSNs enhances alcohol stimulation but not sedation

In two previous studies, we showed that compulsive-like alcohol drinking in mice is associated with changes in *Lrrk2* gene expression in the striatum (*9*) and that mice with enhanced D1R function display higher stimulation and preference for alcohol (*37*). Here we postulated that potentiation of D1R function by deletion of *Lrrk2* from D1-MSN affects alcohol-related behaviors mediated by D1R. To test this, we first assessed the effects of three doses of alcohol (1, 2, and 3 g/kg) on locomotor stimulation. Basal locomotion measured during saline days was similar between genotypes (Fig. 4A; t_(25)_=0.67, *P*=0.51). Alcohol injection induced a dose-dependent and transient increase in locomotion in both genotypes (Fig. 4B), however D1-Lrrk2-KO mice showed higher locomotion at all doses tested, suggestive of a stronger stimulatory response to alcohol (Fig. 4C; REML, genotype effect: F_(1,35)_=4.2, *P*<0.05).

**Figure 4:**
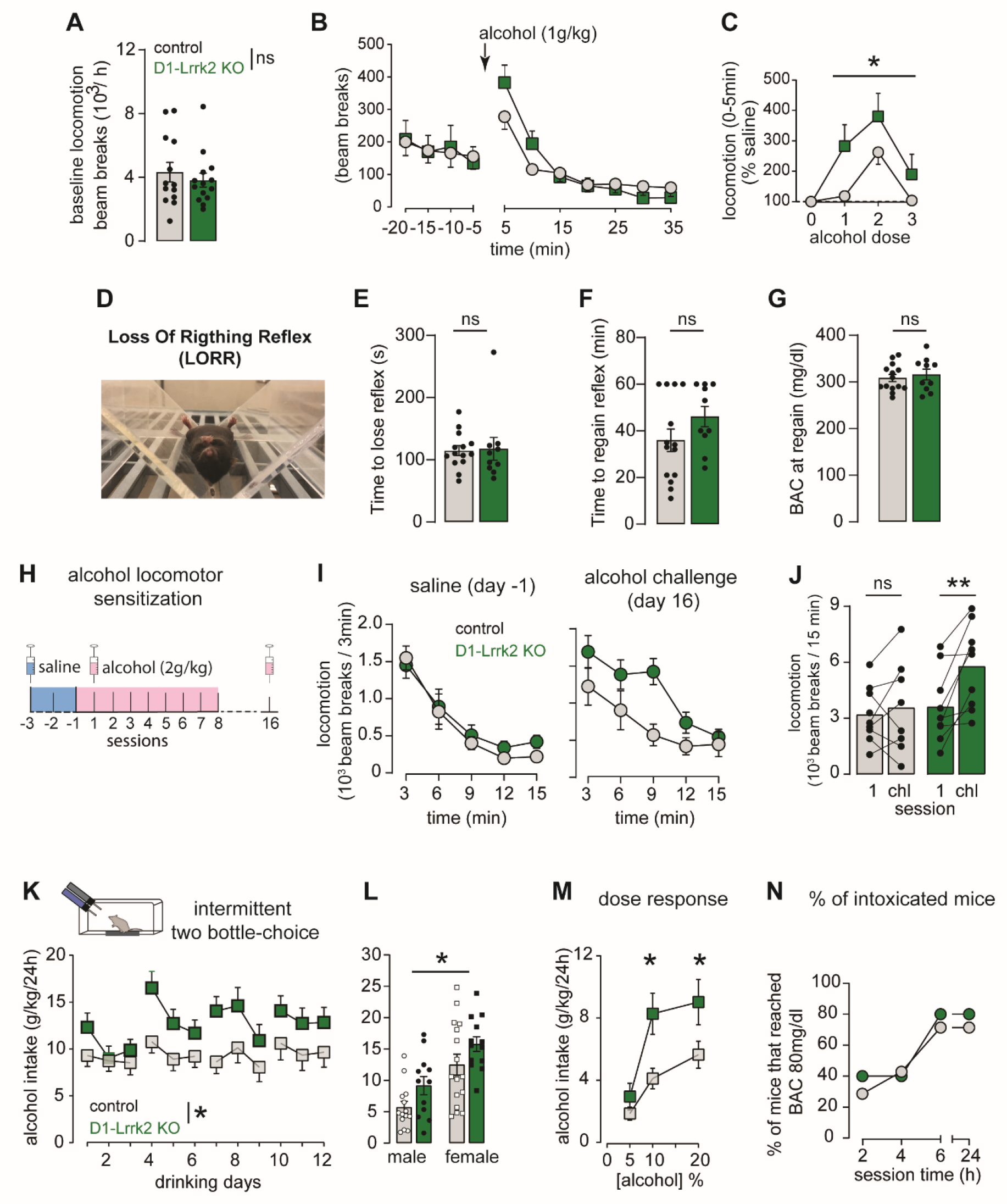
Lrrk2 expression in D1-MSNs regulates alcohol stimulation, locomotor sensitization, and drinking. **A**, Basal locomotion during the habituation session. **B**, Locomotor activity before and after systemic administration of alcohol (1 g/kg). **C**, Dose-dependence of the locomotor response induced by systemic administration of alcohol during the first 5 mins after alcohol injection. **D**, Picture illustrating the LORR task. **E**, Latency to lose the righting reflex (sec). **F**, Time to regain the righting reflex (min). **G**, BAC at the time of regaining the righting reflex. **H**, Graphical representation of the alcohol sensitization protocol. **I**, Basal locomotion during habituation day (saline injection) in alcohol-naive mice *(left)* and during alcohol challenge day (*right*). **J**, Mean locomotor activity during the first day of alcohol injection and the challenge day. **K**, Mean alcohol intake (g/kg/24h) during intermittent two-bottle-choice sessions. **L**, Sex differences in the overall alcohol intake during intermittent two-bottle-choice. **M**, Alcohol intake as a function of the alcohol concentration offered (5,10, 20%). **N**, Percentage of mice that reached blood alcohol intoxication (> 80 mg/dl) at 2, 4, 6, and 24h after the beginning of the alcohol drinking session. For all panels, bars represent mean ± S.E.M and symbols represent values from individual mice. (*) denotes *P* < 0.05; (**) denotes *P*<0.005.

The sedative effects of alcohol were assessed using the loss of righting reflex (LORR) test. Alcohol naïve mice received 3 g/kg (*i*.*p*.) alcohol and the latency to lose and regain the righting reflex (ability to right from a supine position) was recorded (Fig. 4D). Mice from both genotypes showed similar mean latency to LORR (Fig. 4E; t_(22)_=0.16, *P*=0.87), and similar mean time to regain the righting reflex (Fig. 4F; t_(22)_=1.5, *P*=0.15). The blood alcohol concentration (BAC) was measured at the time of regaining the righting reflex and was similar in both genotypes (Fig. 4G; 316 ± 11 *vs* 308 ± 8 mg/dl; t_(22)_=0.56, *P*=0.6). Thus, the enhanced alcohol stimulation is mediated by other mechanisms rather than changes in alcohol metabolism, likely to include enhanced D1R function.

### Loss of LRRK2 function enhances alcohol sensitization and drinking

The development of locomotor sensitization to alcohol is a form of behavioral adaptation dependent on synaptic plasticity mediated by D1R activation in the striatum (*38, 39*). We measured the development and expression of alcohol locomotor sensitization in D1-Lrrk2-KO and control mice following 8 daily injections of alcohol (2g/kg, *i*.*p*.; Fig 4H). Basal locomotor activity was similar between genotypes (Fig. 3I-left; no genotype effect F_(1,16)_=0.26, *P*=0.62). During the challenge test, D1-Lrrk2-KO mice showed a larger locomotor response to alcohol than controls (Fig. 4I-right, 4J). Additionally, locomotor response during the challenge test was larger than on day 1 specifically in D1-Lrrk2-KO mice (Fig. 4J; REML: session effect: F_(1,15)_=9.8, *P*<0.05 and interaction: F_(1,15)_=4.9, *P*<0.05; Sidak’s test, control: *P*=0.78, D1-Lrrk2-KO: *P*<0.005).

Voluntary drinking and preference for alcohol were assessed using an intermittent two-bottle-choice paradigm in which mice had 24h access to alcohol solution (20%) every other day and ad libitum access to water every day for 4 weeks. D1-Lrrk2-KO mice showed a 35% increase in alcohol drinking compared to controls and consumed overall 12.6 ± 5.6 g/kg/day of alcohol while control mice consumed 9.3 ± 1.2 (Fig. 4K; genotype effect: F_(1,53)_=4.0, *P*<0.05). The increase in drinking was observed in both female and male mice (Fig. 4L; genotype effect: F_(1,51)_=6.0; *P*<0.05). D1-Lrrk2-KO mice also showed higher alcohol preference compared to control mice (33.3 ± 3.9% *vs* 46.6% ± 19%; t_(53)_=2.45, *P*<0.05).

An independent cohort of alcohol naïve mice was used to assess dose dependency of alcohol drinking. Compared to controls, D1-Lrrk2-KO mice consumed more alcohol when given access to 10% and 20% alcohol solution (Fig. 4M; genotype effect: F_(1,21)_=5.5, *P*<0.05; Tukey’s test, *P*<0.05). Blood was collected at different time points in a subset of mice during the 2-bottle choice experiment and BAC was measured. By 2h after the start of the session, 40% of D1-Lrrk2-KO mice and 28% of controls reached alcohol intoxication levels in the blood (>80 mg/dl). By 24h, 80% of D1-Lrrk2-KO and 71% of controls reached 80 mg/dl at some point during the 24h drinking period (Fig. 4N), suggesting that mice consumed alcohol because of its pharmacological effects.

Since the EY262-cre line was used to generate the D1-Lrrk2-KO mouse line, we assessed alcohol consumption in EY262-cre positive and EY262-cre negative littermate controls. During 8 sessions with access to 20% alcohol solution, both genotypes consumed similar amounts of alcohol (Fig. S2A; no genotype effect: F_(1,14)_=0.46, *P*=0.51), ruling out possible off-target effects of Cre expression in D1R positive cells. Finally, body weight and water consumption were similar for both genotypes, suggesting the differences seen in D1-Lrrk2-KO mice are not due to altered liquid consumption (Fig. S2B; t_(24)_=0.83, *P*=0.42) or differences in body weight (Fig. S2C-D; females: t_(46)_=1.14; males: t_(37)_=0.17, *Ps*>0.05).

### Enhanced alcohol response is selective for mice lacking *Lrrk2* in D1-MSNs

To test whether changes in alcohol-related behaviors were contingent on the deletion of the *Lrrk2* gene specifically from D1-MSNs, we tested alcohol-induced stimulation and voluntarily alcohol consumption in mice lacking the *Lrrk2* gene selectively in D2-MSN. Lrrk2^loxP/loxP^ mice were crossed with Adora2a-cre mice, which express Cre in D2-MSNs within the striatum (*40-42*). qPCR analysis showed 32% reduction of *Lrrk2* mRNA levels in striatal samples from A2a-Lrrk2-KO and no change in lung tissue (Fig 5A; t_(5)_=4, *P*<0.05; t_(5)_=1.4, *P*=0.2).

**Figure 5:**
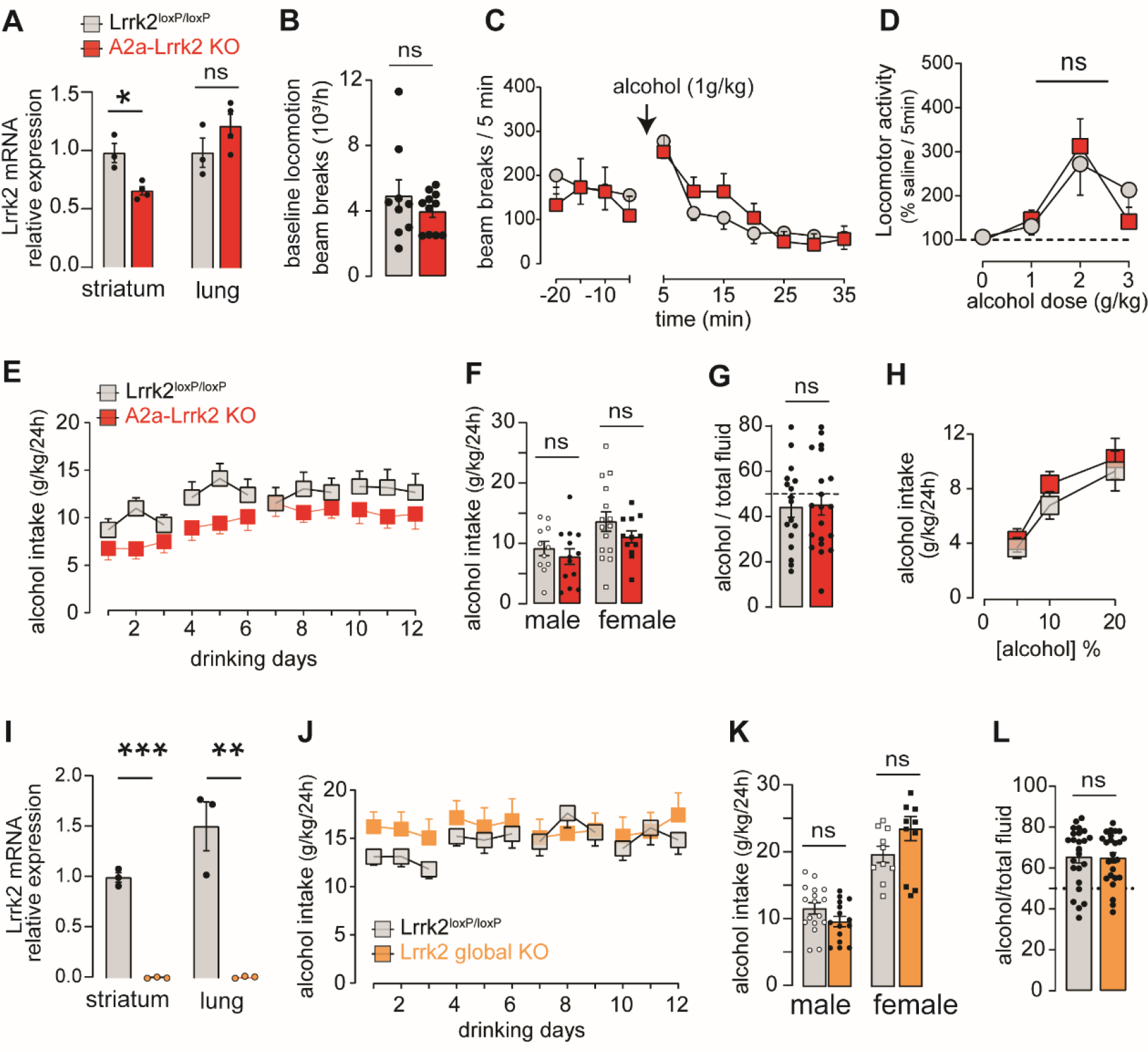
Enhancement of alcohol drinking and alcohol stimulation is selectively modulated by Lrrk2 expression in D1-MSN. **A**, Levels of *Lrrk2* mRNA expression in the striatum and lung tissue of A2a-Lrrk2-KO mice (red) and littermate controls (gray). **B-C**, Basal locomotion measured during habituation session (B) and systemic administration of alcohol (C). **D**, Dose-dependence of the locomotor response induced by systemic administration of alcohol. **E**, Mean alcohol intake (g/kg/24h) during intermittent two-bottle-choice sessions. **F**, Sex differences in the overall alcohol intake. **G**, Alcohol preference. **H**, Alcohol intake as a function of the alcohol concentration. **I**, Levels of *Lrrk2* mRNA expression in the striatum and lung tissue of Global-Lrrk2-KO mice (orange) and littermate controls (gray). **J**, Mean alcohol intake (g/kg/24h) during intermittent two-bottle-choice sessions. **K**, Sex differences in the overall alcohol intake. **L**, Alcohol preference. For all panels, bars represent mean ± S.E.M and symbols represent values from individual mice. (*) denotes *P* < 0.05; (**) denotes P<0.01; (***) denotes *P*<0.001.

Basal locomotion during saline days was similar between genotypes (Fig. 5B; t_(18)_=0.97, *P*=0.34). Alcohol produced a transient dose-dependent increase in locomotion that was similar in A2a-Lrrk2-KO and control mice at all 3 doses tested (Fig. 5C-D; REML: no genotype effect: F_(1,31)_=0.02, *P*=0.9). Voluntary alcohol drinking (Fig. 5E-F; F_(1,49)_=2.6, *P*=0.11) and preference (Fig. 5G; t_(35)_=0.17; *P*=0.86) was similar between A2a-Lrrk2-KO and controls. No difference was found when alcohol consumption was measured in a dose-response fashion using a separate cohort of naïve mice. (Fig. 5H; no genotype effect: F_(1,31)_=0.45, *P*=0.5).

Last, we assessed the effect of global deletion of the *Lrrk2* gene on alcohol consumption. qPCR analysis showed a total loss of *Lrrk2* mRNA in both striatum and lung tissue samples from Global-Lrrk2-KO, confirming the global deletion of the *Lrrk2* gene (Fig. 5I; t_(4)_=20, *P*<0.0001; t_(4)_=6.2, *P*<0.005). Global-Lrrk2-KO mice consumed similar amounts of alcohol than controls (Fig. 5J-K; no genotype effect: REML, F_(1,52)_=0.41, *P*=0.52) and showed similar alcohol preference (Fig. 5L; t_(47)_=0.14, *P*=0.9). Water consumption (Fig. S2B; t_(10)_=1.5, p=0.17; t_(8)_=0.67, p=0.52) and body weight were also similar across genotypes (Fig. S2C; males: t_(36)_=0.93 and t_(33)_=0.16; females: t_(33)_=0 and t_(20)_=0.2, *Ps*>0.05; A2a-Lrrk2 and Global-KO, respectively). Thus, increased alcohol response and drinking appear selective to mice with *Lrrk2* deletion selectively from D1-MSNs.

### Alcohol lowers LRRK2 kinase activity in the dorsal striatum

Using western blot analysis in C57BL6/J wild-type mice we tested whether alcohol drinking modulates LRRK2 kinase activity in the striatum. LRRK2 activity was assessed via quantification of the phosphorylation levels of LRRK2 at residue S935, a phospho-site that is decreased by LRRK2 kinase inhibitors, and via phosphorylation levels of the LRRK2 substrate Rab10, a small GTPase that is phosphorylated by LRRK2 at residue T73 (*43, 44*). C57Bl6/J mice were given free access to 20% alcohol and water for 24 hours and LRRK2 activity was measured in the DMS and dorsolateral striatum (DLS) 48h after the alcohol drinking session (Fig. 6A). Alcohol consumption promoted a general decrease in the levels of pS935-LRRK2 in both striatal subregions (Fig. 6B; alcohol effect: F_(1,18)_=5.3, *P*<0.05; no interaction: F_(1,18)_=0.4, *P*=0.53) and a specific decrease in the levels of pT73-Rab10 in the DMS (Fig. 6C; interaction: F_(1,17)_=14.4 *P*<0.005; DMS Sidak’s test *P*<0.05). Furthermore, the reduction of pT73-Rab10 was correlated with the amount of alcohol consumed (Fig. 6D; *r*^2^ = 0.33, *P*<0.01). Note that the correlation is significant even after removing the animal with high alcohol intake (>21 g/kg) from the dataset (*r*^*2*^*=*0.23; *p*<0.05). Total levels of LRRK2 and Rab10 were not affected by alcohol drinking (Fig. S3B; no genotype effect: F_(1,18)_=0.56; F_(1,18)_=0, *P*=0.99). However, there was a regional difference in the levels of LRRK2, which were higher in the DLS compared to the DMS (Fig. S3B; region effect: F_(1,18)_=42, *P*<0.0001).

**Figure 6:**
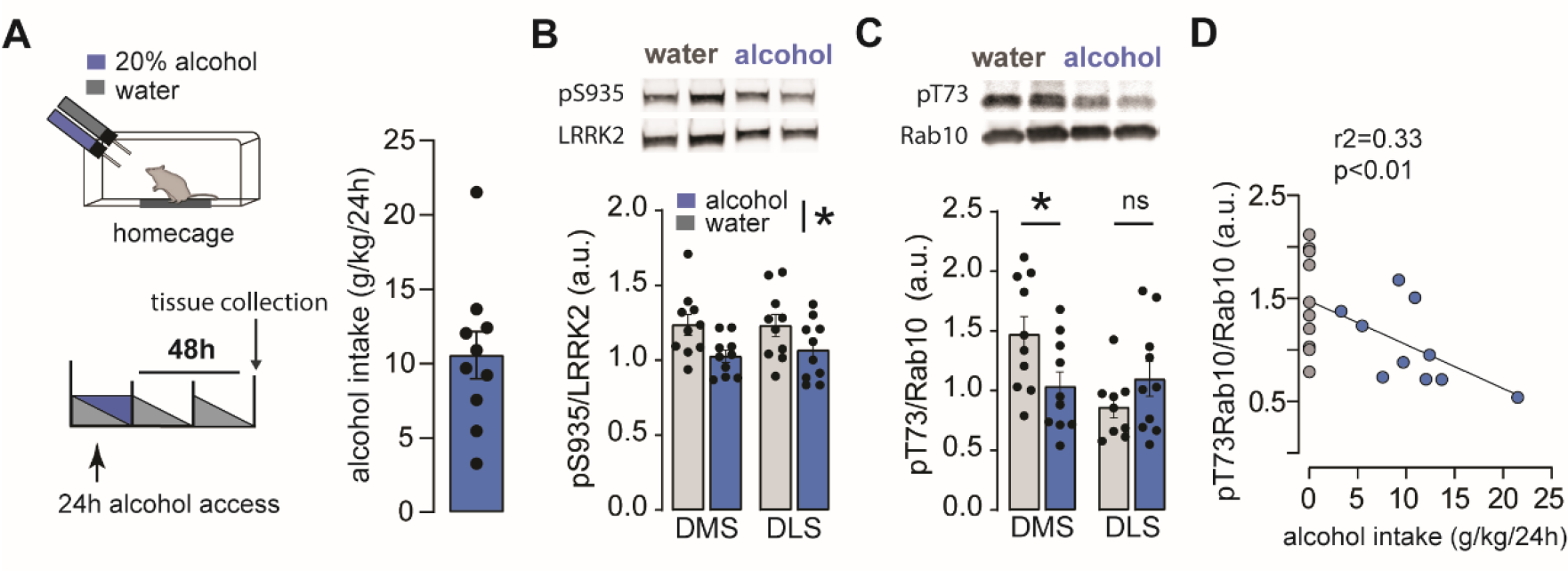
Alcohol modulates LRRK2 activity in mouse striatum. **A**, *Top*, schematic diagram of the two-bottle-choice paradigm used to measure volitional alcohol drinking over 24h. *Bottom*, time course of the experiment outlines 24h alcohol drinking and brain tissue collection for western blots (WB) 48h after the single drinking session. R*ight*, Average alcohol consumed over 24h. **B-C**, *Top*, Images of Western Blots for pS935 LRRK2 and total LRRK2 (*B*) and pT73-Rab10 and total Rab10 (*C*) in samples from DMS and DLS of water and alcohol drinking mice. *Bottom*, bar graphs showing the ratio of labeling density for the phosphorylated forms of LRRK2 and Rab10 over the total protein levels. **D**, pT73-Rab10 as a function of the amount of alcohol consumed over a single 24h drinking session. For all panels, bars represent mean ± S.E.M and symbols represent values from individual mice. * denotes *P* < 0.05.

Passive administration of alcohol (2 g/kg, *i*.*p*.) produced a significant reduction in pT73-Rab10 levels which peaked after 12h of administration and was recovered by 48 h (Fig. S3D; time effect: F_(3,12)_=5.2, *P*<0.05; Tukey’s test: Saline *vs* 12h, *P*<0.05). No significant change in pS935-LRRK2 levels was observed, but there was a trend of reduction (Fig. S3E; effect of time: F_(3,12)_=2.7, *P*=0.09). Note that this passive alcohol administration increased BAC to levels exceeding 80 mg/dl, the legal limit considered for intoxication, for several hours (Fig. S3D). Thus, both voluntary drinking and passive administration of alcohol decreases LRRK2 kinase activity in the striatum, which is suggestive of a bidirectional regulation between LRRK2 activity and alcohol.

### Enhanced likelihood of heavy and punishment-resistant alcohol drinking

Heightened motivation to consume alcohol and persistent drinking despite negative consequences are two core manifestations of AUD (*45, 46*). Here, we used an operant alcohol self-administration paradigm (SA) to assess motivation and resistance to punishment (Fig. S4A). Before the operant training, mice were first exposed to a drinking in the dark (DID) procedure for 4 weeks. Average alcohol intake during DID was similar between genotypes (3.9 ± 0.2 g/kg/4h vs 4.0 ± 0.3 g/kg/4h; t_(26)_=0.23, *P*>0.05). Mice were then trained to press an active lever to gain access to a sipper tube containing 20% alcohol solution for 60s. D1-Lrrk2-KO mice showed a higher rate of responding on the active lever and earned longer access time to alcohol compared to controls (Fig. 7A-B; genotype: F_(1,25)_=5.7; session: F_(3.5,88)_=15 and interaction: F_(12,300)_=2.6, *Ps*<0.05). Mice from both genotypes learned to discriminate between active and inactive levers (Fig. S4B; lever response: F_(1,14)_=16, *P*<0.005). D1-Lrrk2-KO mice showed a strong trend towards larger alcohol consumption during the 6h access than controls (Fig. 7C; 3.3 ± 0.6 g/kg *vs* 1.8 ± 0.4 g/kg; F_(1,25)_=4, *P*=0.056). Overall, the cumulative alcohol consumption of D1-Lrrk2-KO mice was 1.8 fold significantly higher than littermate controls by the end of the SA training sessions (Fig. 7D; 23.9 ± 5.6 vs 43.5 ± 29.3g/kg; interaction: F_(12,300)_=4.22; *P*<0.0001). The rate of licking in the sipper spout was similar across genotypes (Fig. S4C, no genotype effect: F_(1,25)_=1.38, *P*=0.21). Interestingly, 40% of control mice showed minimal or no alcohol intake (<0.1 g/kg/6-hour) and failed to reach the acquisition criterion for the operant task. This percentage was significantly larger compared to D1-Lrrk2-KO mice where only 17% failed to reach the criteria (Fig. 7E).

**Figure 7:**
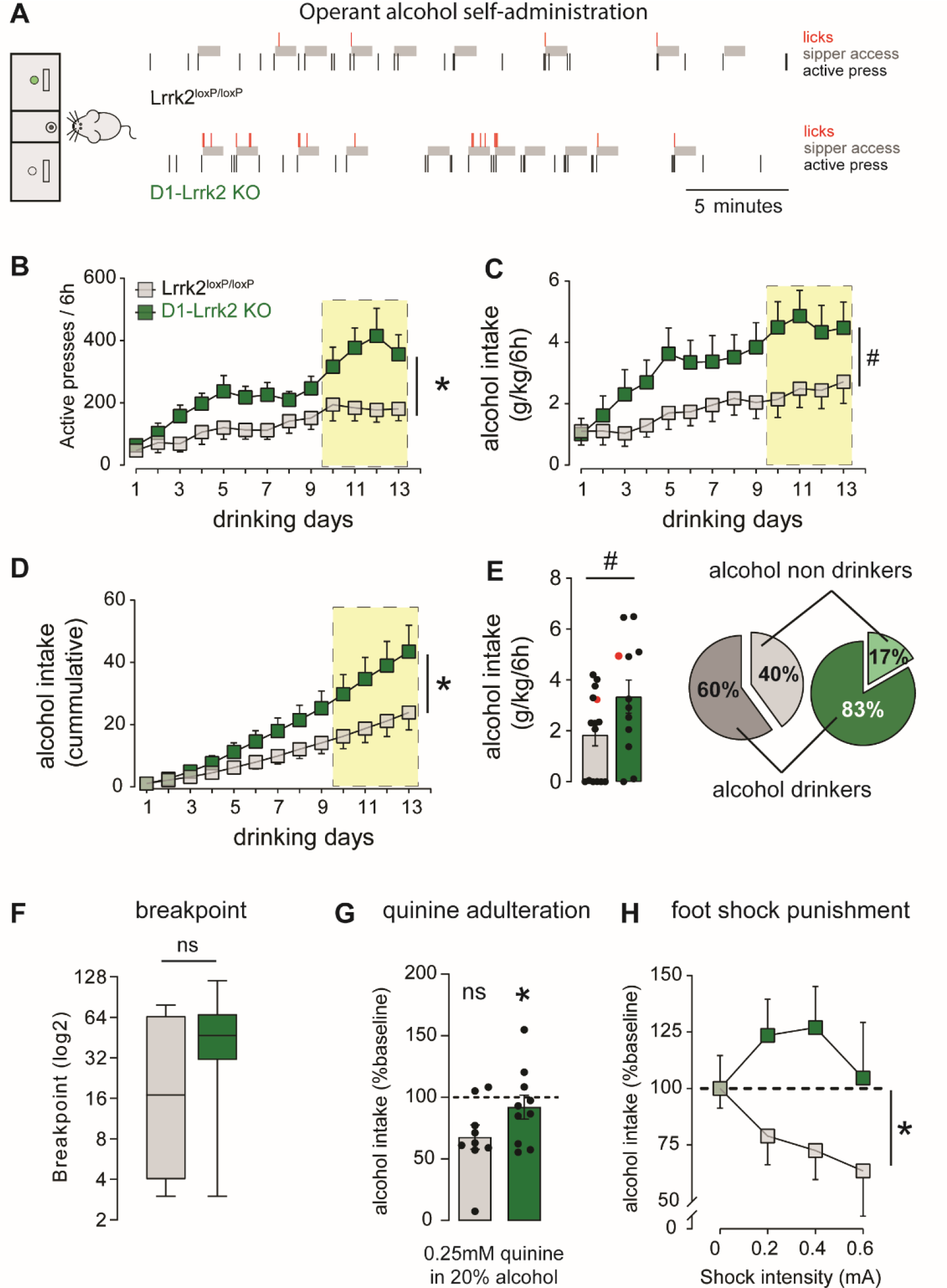
LRRK2 expression in D1R-expressing neurons promotes resilience to compulsive alcohol drinking in mice. **A**, Representative raster plot of the first 30 min of a training session at FR3 for a control mouse (top) and a D1-Lrrk2-KO mouse (bottom). Three responses in the active lever (black bar) lead to 60 seconds of access to a sipper tube containing 20% alcohol (gray shaded area). Licks responses are shown in red. **B**, Mean rate of active lever pressing during the operant drinking sessions. Shaded yellow area represents FR3 sessions. **C**, Average daily alcohol intake (g/kg/6h). **D**, Average cumulative daily alcohol intake (g/kg/6h). **E**, *(left)* Average of alcohol drinking (g/kg/6h). Red dots represent mice used as examples in figure 7A. (*Right)* Proportion of alcohol drinkers and non-drinkers. **F**, Breakpoint during progressive ratio session. **G-H**, Percent of change in alcohol intake during a quinine adulteration session (G) and during foot shock sessions (H) compared to baseline consumption. For all panels, bars represent mean ± S.E.M and symbols represent values from individual mice. * denotes *P* < 0.05; (#) denotes *P*=0.056.

Mice that reached the criteria for acquisition were further tested in a single progressive ratio (PR) session and two different kinds of punishment sessions used to assess persistence of drinking despite negative consequences: quinine adulteration (0.25 mM) and foot-shock punishment (Fig. S4A). The median breakpoint was similar between D1-Lrrk2-KO mice and controls (Fig. 7F; Mann-Whitney U_(30)_; *P*=0.23). During the quinine adulteration session, control mice showed a significant reduction in alcohol drinking while D1-Lrrk2-KO mice maintained an average intake similar to baseline levels (Fig. 7G; one-sample t-test, D1-Lrrk2-KO: t_(8)_=3.3, *P*<0.05; control: t_(9)_=0.8, *P*=0.44). Neither genotype showed a significant change in lever press rate, which suggests that taste adulteration mainly suppressed drinking behavior but not seeking behavior (Fig. S4D, right; no effect of dose F_(1,17)_=0.22, *P*=0.65). We also assessed the suppression of alcohol drinking by a higher concentration of quinine using a non-operant paradigm (two-bottle choice) in a separated cohort of mice. Both genotypes dramatically reduced drinking when alcohol was adulterated with 0.5 mM quinine, however, D1-Lrrk2-KO mice showed only a small reduction in drinking when alcohol was adulterated with 0.25 mM quinine (Fig. S4E; genotype effect: F_(1,19)_=5.0, *P*<0.05; Sidak’s test, *P*<0.05). No genotypic differences in taste sensitivity to quinine were found when a preference test was assessed between tap water and water adulterated with 0.25 and 0.5mM quinine. Both genotypes showed a very low preference for quinine solution (0-6%), indicating no alteration in bitter taste sensitivity and suggesting that the differences in adulterated alcohol drinking are driven by changes in the response to alcohol (Fig. S4F; no genotype effect F_(1,12)_=1.4, *P*=0.25).

During foot shock sessions, D1-Lrrk2-KO mice maintained high levels of alcohol drinking at all shock intensities whereas controls substantially reduced alcohol consumption (Fig. 7H; genotype effect: F_(1,17)_=5.6, *P*<0.05). D1-Lrrk2-KO mice, but not controls, showed a reduction in the number of lever presses that was dependent on shock intensity (Fig. S3G), however, despite the reduction in the number of lever presses, D1-Lrrk2-KO mice earned more rewards than littermate control during foot shock sessions (Fig. S4H).

Two sets of important control experiments were carried out to assess the validity of the punishment-induced reduction of alcohol drinking. First, we measured pain threshold sensitivity in D1-Lrrk2-KO using the hot plate test and found similar latencies to withdraw and lick the forepaw than control mice (Fig. S4I; t_(12)_=0.51, *P*=0.6). Second, we assessed the effect of quinine adulteration and foot shock on sucrose SA behavior. Both genotypes self-administered similar amounts of sucrose at all concentrations tested (Fig. S5A-B; no genotype effect: F_(1,44)_=1.4, *P*=0.24; t_(44)_=1.1, *P*=0.26). Sixty-five percent of control mice and 48% of D1-Lrrk2-KO mice did not consume sucrose during the task and failed to reach acquisition criterion for the operant task (non-learners) (Fig. S5C; *X*^*2*^_(1,45)_=0.75, *P*=0.38). D1-Lrrk2-KO mice and control showed similar breakpoints during the progressive responding session (Fig. S5D; U_(23)_, *P*=0.23). Adulteration of sucrose solution with 0.25 mM quinine produced a robust suppression in drinking in both genotypes (Fig. S5E; t_(16)_=0.17, *P*=0.86). Both genotypes also reduced during the foot shock punishment session, 93% reduction in control mice and 85% reduction in D1-Lrrk2-KO (Fig. S5E; t_(16)_=0.73, *P*=0.48).

Altogether, the findings of this study provide strong evidence for a novel role of LRRK2 function in regulating D1R signaling in striatal D1-MSNs and it shows that impair LRRK2 function in D1R-expressing neurons, but not D2R-expressing neurons, leads to significant alterations in striatal circuitry that promote alcohol seeking and alcohol drinking despite adverse consequences.

## Discussion

This study uncovers a novel role of the Parkinson’s-related gene *Lrrk2* in regulating D1R function and alcohol drinking. Taking advantage of conditional *Lrrk2* mice to generate a cell-specific knockout, we dissected the contributions of LRRK2 to dopamine-related behaviors and the cellular and behavioral response to alcohol. We identified that loss of LRRK2 function in striatal direct-pathway neurons potentiates D1R signaling and function, enhances the reinforcing properties of alcohol, and promotes the development of punishment-resistant compulsive-like alcohol drinking. Two previous studies have shown associations between *Lrrk2* expression in the brain and alcohol behaviors in rodents and zebrafish but the mechanism underlying the association was unknown (*9, 10*). This study provides a mechanistic understanding of how low levels of LRRK2 in D1-MSNs lead to higher alcohol drinking and punishment-resistant drinking behavior in mice.

LRRK2 is a complex multidomain protein involved in several important cell signaling pathways, such as Wnt-, MAPK-, and PKA-signaling. These intracellular pathways modulate a broad range of neuronal functions, including vesicular trafficking, autophagy, and cytoskeleton dynamic (*47-52*). Mutations in human *LRRK2* are a common genetic cause of Parkinson’s disease indicating that LRRK2 has an important role in the regulation of basal ganglia function (*8*). Further, a variety of non-motor abnormalities have been associated with dysfunction of the *Lrrk2* gene in rodents; from anxiety-like and depressive-like behaviors to alterations in the stress response and social interaction (*53-56*). Here we found that deletion of *Lrrk2* from D1-MSN promoted potentiation of D1R function, which was observed at multiple levels. At the synaptic level, we observed a postsynaptic effect at the soma of D1-MSNs where D1R-like agonist increased neuronal excitability and a presynaptic effect in the axon terminals of D1-MSNs in the SNr where D1R-like agonist potentiated synaptic transmission in SNr neurons. These effects were not observed in wild-type littermate control mice, therefore loss of the *Lrrk2* gene in these neurons seems to promote a gain of function, maybe via a sensitized-like state of these receptors, as described in rodents following dopamine depletion (*57, 58*). We proposed then a physiological role for LRRK2 in D1-MSNs to limit D1R signaling at both cellular compartments: soma, and axon terminals. These findings stand in support of previous studies which indicated that LRRK2 modulates PKA signaling downstream of D1R (*19*) and regulates the receptor internalization (*20, 21*).

At the cellular level, we found that constitutive cell-specific deletion of *Lrrk2* increases the D1R-like agonist mediated induction of immediate-early gene c-Fos expression in D1-MSNs. However, postnatal deletion of the *Lrrk2* gene in fully developed brains did not reproduce the effect. This result suggests that loss of LRRK2 function early in development is required for D1R potentiation or that cell-specificity in the deletion is important. Due to lack of cre-expressing viral vectors with specific promoter for D1-MSNs, we were unable to delete of *Lrrk2* gene in a cell-specific manner in postnatal brains in this study.

At the behavioral level, deletion of the *Lrrk2* gene from D1-MSNs did not change baseline levels of dopamine-related behaviors such as locomotion, velocity, and response to novelty but it rather specifically enhanced alcohol-related behaviors that have been known to be dependent on D1R activity, such as alcohol stimulation, sensitization, and intake. Interestingly, loss of LRRK2 function in D1-MSN did not affect alcohol-induced sedation, which is in agreement with previous findings suggesting that alcohol sedation is unlikely to be modulated by or dependent on dopamine receptor activity or dopamine signaling (*59-61*).

Interestingly, we recently showed that mice that drink high levels of alcohol are more sensitive to the stimulatory effects of alcohol compared to low drinkers (*37*). Both alcohol-induced stimulation and alcohol drinking required activation of D1R in the dorsomedial striatum (*23, 37, 62, 63*). Individuals who display high stimulation and low sedation to alcohol are at higher risk of becoming heavy drinkers and struggle in controlling consumption. This association was first reported in humans, and these individuals are more prone to alcohol abuse. (*3, 37, 64-66*). This link appears conserved in rodents, where higher alcohol stimulation is predictive of higher alcohol preference, escalation, and higher resistance to punishment when paired with alcohol (*37*). Additionally, D1-MSN activity modulates alcohol rewarding properties and alcohol drinking. Chemogenetic stimulation of D1-MSN in the dorsal striatum increases alcohol drinking while chemogenetic or pharmacological inhibition decreases alcohol drinking in mice (*62, 67*). Here, we found that D1-Lrrk2-KO mice show higher alcohol preference consumption than controls. Importantly, mice of both genotypes reached blood alcohol levels above the intoxication level of 80 mg/dl during intermittent alcohol access suggesting that alcohol was consumed because of its pharmacological effects and that the alcohol consumption reflects differences in reinforcing properties of alcohol.

Loss of control over alcohol drinking is a hallmark of AUD which is reflected as high motivation to consume alcohol and continued drinking despite negative consequences (*46, 68*). Therefore, in this study, we also used an operant paradigm of alcohol self-administration which allows for measurements of motivation to seek and the persistence of the behavior even when paired with punishment (*69-71*). D1-Lrrk2-KO mice displayed a higher rate of responding on the active lever that granted access to alcohol and showed overall higher levels of alcohol consumption than controls. D1-Lrrk2-KO also were more likely to continue drinking despite negative consequences, as measured during punishment sessions using both quinine-adulteration and foot-shock pairing. Pain sensitivity was not altered in the D1R-specific KO mice nor was the ability of quinine to suppress water drinking. Thus, differences in pain threshold and taste sensitivity are not responsible for driving the punishment-resistant alcohol drinking in D1-Lrrk2-KO mice. Interestingly, there was no difference in breakpoint response between D1-Lrrk2-KO mice and controls, suggesting that deletion of *Lrrk2* does not affect motivation to drink or seek alcohol. Additionally, while operant responses during quinine adulteration and punishment sessions were highly correlated, breakpoint responses were not. This dissociation between motivational salience and response to punishment is in line with the literature on pre-clinical models of AUD (*72, 73*).

Importantly, increased alcohol consumption is not justified by a generalized increase in motivation for natural rewards or taste sensitivity because 1) there was no general increase in sucrose preference or operant sucrose drinking; 2) foot shock punishment and quinine adulteration promoted a robust suppression in sucrose consumption but not in alcohol drinking; 3) there was no difference in the preference for non-caloric sweetener; 4) there was no difference in regular food or high-fat diet consumption.

Notably, the changes in alcohol-related behaviors are only observed in mice lacking *Lrrk2* in D1-MSNs but not in those lacking the gene in D2-MSNs or when the *Lrrk2* deletion is global. This remarkable cell-specificity resembles the findings observed with DARPP-32, which led to different outcomes following global or pathway-specific deletion in the striatum (*74, 75*). In the case of DARPP-32, it was proposed that global deletion does not lead to locomotor deficits because deletion in one pathway (D1-MSNs for example) counteracts the effect of deletion on the other pathway (D2-MSNs). The behavioral output is thought to be dependent rather on the balance of activity between D1-MSNs and D2-MSNs; thus, deletion in both cell types can lead to no net change when the effects are counterbalanced. While the loss of *Lrrk2* in D2-MSNs alone was not sufficient to change alcohol drinking significantly, it appears sufficient to prevent the potentiation of D1R function and drinking when *Lrrk2* is deleted in both cell types. Thus, one could speculate that the lack of phenotype in the Global knockout mice is evidence that the balance activity of the two pathways (D1-MSNs and D2-MSNs) contributes to control over alcohol drinking. However, it is worth noting that *Lrrk2* is expressed in many other brain regions as well as different cell types, and therefore the lack of phenotype in the Global-Lrrk2-KO mice could be the result of a more complex phenomenon. Another recent study by Jadhav and colleagues supports the concept of balance of activity. Using a similar self-administration approach to assess the compulsive aspect of alcohol drinking, the authors found that rats that develop uncontrolled alcohol-seeking have higher levels of *Drd1a* mRNA and lower levels of *Drd2* mRNA in the striatum (*76*).

Our study reveals multiple alterations within the striatum upon loss of LRRK2 function in D1-MSNs. However, we speculate that the mechanism driving vulnerability to alcohol abuse extends beyond the striatum. In fact, the knockout strategy used in this study promotes the deletion of the *Lrrk2* gene not only in D1-MSNs in the striatum, but rather in all D1R-expressing neurons, which include neurons from different cortical areas, amygdala, and hippocampus. It is also possible that developmental alterations are in play because of the constitutive deletion of *Lrrk2*. Further studies investigating the consequences of postnatal disruption of the *Lrrk2* gene or pharmacological inhibition of LRRK2 kinase activity will help in assessing the developmental role.

Finally, we found that a single episode of heavy alcohol drinking was sufficient to decrease LRRK2 kinase activity. Phosphorylation of the LRRK2 kinase substrate Rab10, a molecule involved in the regulation of protein transport to the plasma membrane, was significantly decreased after 24-hour alcohol drinking. These pieces of biochemical evidence suggest that the regulation of alcohol drinking by LRRK2 function is bidirectional, and that alcohol drinking can also modulate LRRK2 kinase activity.

Based on all these findings, we propose that an important function of LRRK2 in striatal D1-MSNs is to act as a negative regulator of D1Rs. LRRK2 then limits D1R signaling in the striatum and lowers alcohol’s reinforcing properties, promoting resilience to compulsive alcohol drinking. Pharmacological manipulations that modulate LRRK2 function might hold therapeutic promise in reducing heavy uncontrolled drinking of alcohol.

## Materials and Methods

### Animals

All experiments were approved and performed in accordance with guidelines from the National Institute on Alcohol Abuse and Alcoholism’s Animal Care and Use Committee. Male and female mice (8-18-week-old) were used in all experiments. Data from both sexes were collapsed if no sex differences were found. Lrrk2^loxP/loxP^ mice were generated (*77*) and generously provided by Dr. Cai (NIA, NIH). Two different lines of cell-type specific Lrrk2 knockout mice were generated by crossing Lrrk2^loxP/loxP^ mice with Drd1a-cre (B6.FVB(Cg)-Tg(Drd1-cre)EY262Gsat/Mmucd) and Adora2a-cre (B6.FVB(CG)-TG(ADORA2A-CRE)KG139GSAT/MMUCD) mouse lines. The Global-Lrrk2-KO mouse (B6.129X1(FVB)-LRRK2TM1.1CAI/J) was used for global deletion of the *Lrrk2* gene. All transgenic mouse lines were backcrossed to C57BL6/J background for several generations. A D1Td tomato reporter line (B6.Cg-Tg(Drd1a-tdTomato)6Calak/J) was used for specific experiments as detailed below. For all experiments using transgenic mice, Cre negative Lrrk2^loxP/loxP^ littermates were used as controls. All mice were genotyped at weaning using real-time PCR with their respective probes by Transnetyx (Cordova, TN). Mice used in experiments that did not require alcohol drinking were grouped-housed under normal light cycle (12h dark/light, lights on 6:30 am). For alcohol drinking experiments, mice were singled-housed and transferred to reverse light cycle (12h dark/light, lights off 6:30 am) at least 10 days before the start of the experiments. Standard rodent chow and water were always available *ad libitum* in home cage, except during DID sessions when water was removed for 4 hours/session (no food deprivation was used in any experiment).

### Lrrk2 mRNA expression analysis

FASTQ files were downloaded from GEO and ENA datasets (GSE81672: Nac, PFC, BLA, vHipp; GSE89692: PFC, VTA and PRJEB36194: DMS) using SRA toolkit. Only naïve C57BL6/J wild type mice were used in this analysis. Read quality, length, and composition were assessed using FastQC before trimming low quality bases (Phred < 20) and adapters using Fastp. Alignment to the GRCm39 Gencode genome assembly and gene-level counts were generated using STAR with default options. BiomaRt in the R (version 4.0.3) was used to calculate FPKM of gene counts. Analyses were performed using the NIH HPC Biowulf cluster (http://hpc.nih.gov).

### Tissue and cell-specific quantification of Lrrk2 Mrna

For quantitative real time PCR (qPCR), tissue samples from dorsal striatum and lung were collected, quickly frozen in liquid nitrogen and stored at -80°C until RNA extraction. RNA was extracted using RNAeasy mini kit (Qiagen) according to manufacturer instructions. RNA samples were treated with DNAse and quantified on spectrophotometer (DeNovix, USA). RNA quality was assessed via spectrometry (260/280 ratio ≥ 1.9; 230/260 ratios ≥ 1.7). Total RNA (100 ng) was converted to cDNA using iScript (BioRad) according manufacture protocol. qPCR was performed using TaqMan Fast Advanced Master Mix (Thermo Fisher Scientific) using probes for Lrrk2 (Mm01304130_m1) and Gapdh (Mm99999915_g1). Relative quantification of Lrrk2 transcript was calculated according ΔΔCt method and normalized by Gapdh expression and by littermate control levels. For RNAscope, reagents, probes and equipment from Advanced Cell Diagnostics were used. Coronal sections (16μm) were sliced on cryostat at -20°C. Brain slices containing striatum (∼AP: +1.1 mm from Bregma) were mounted on Superfrost slides (Thermo Fisher Scientific) and processed as follow: post-fixed in 4% paraformaldehyde for 30 min; dehydrated in increasing ethanol solutions (50%, 70%, 100%; 5 min each); incubated with protease IV (30 min at RT); washed; and hybridized with probe set (C1-Lrrk2, C2-Drd1 and C3-Adora2a) for 2 hours at 40°C. Slides were washed; incubated with Amp1-4C reagents; and mounted using ProLong™ Gold Antifade (Thermo Fisher Scientific).

### LRRK2 Immunostaining

Protocol was adapted from Davies and colleagues (*78*) and is briefly described below. Naive D1-Lrrk2-KO and littermate controls expressing tdTomato under the Drd1a promoter (4 mice/group, 3-4 slices per animal) were anesthetized and transcardially perfused with 50 ml of ice-cold PBS containing 0.025% heparin followed by 4% PFA (paraformaldehyde). Brains were dissected, post-fixed for 2 h in 4% PFA at 4°C and immediately sectioned in coronal slices (50 μm) using a vibratome (Ted Pella). Coronal sections were incubated with 10 mM sodium citrate (pH 6.0) containing 0.05% Tween 20 for 30 min at 37°C with agitation to enable antigen retrieval followed by three 5-min washes in TBS. Non-specific sites were blocked by incubating sections in 5% Goat Normal Serum in TBS containing 0.3% Triton X-100 for 1 h at 4°C with agitation. Next, sections were incubated in primary antibody [MJFF2 (c41-2)] (1:50, Abcam, #ab133474) diluted in 5% Goat Normal Serum in TBS containing 0.15% Triton X-100 for 48 h at 4°C with agitation. ‘No primary’ antibody and concentration-matched IgG controls were included in each experiment. Sections were then incubated in secondary antibody (Alexa Fluor® 488, Abcam, #ab150077, 1:1000) 18 h at 4°C with agitation. Sections were mounted with Vectashield with DAPI (Vector Laboratories).

### c-Fos Immunostaining

D1-Lrrk2-KO and littermate control mice expressing tdTomato under the Drd1a promoter were injected with either SKF 81297 (2mg/kg) or saline (10 ml/kg) 90 mins prior to transcardial perfusion with 4% PFA. Brains were removed and post-fixed in 4% PFA overnight at 4 °C. Coronal sections (50 μm) from each treatment and genotype groups were processed in parallel using a vibratome (Ted Pella). Sections were washed in PBS (3 × 10 min) and blocked with 10% normal goat serum for 2 h at RT. Sections were incubated in primary antibody rabbit mAb cFos (1:500, Cell Signaling Technology, #2250S) for 48 h at 4 °C, washed in PBS (3 × 10 min), then incubated in the secondary antibody anti-rabbit Alexa Fluor 488 (1:500, Invitrogen, #A11008) or Goat anti-Rabbit IgG (H+L) Cross-Adsorbed Secondary Antibody, Cyanine5 (1:500, ThermoFisher, #A10523) for 2 h at RT and washed with PBS (1 × 10 min), and 0.1 M PB (1 × 10 min). Slices from the microinjection experiments were additionally incubated with primary antibody Chicken IgY GFP Polyclonal Antibody (1:100, ThermoFisher, #A10262) for 48 h at 4 °C and with secondary antibody Goat anti-Chicken IgY (H+L) Cross-Adsorbed Secondary Antibody, Alexa Fluor Plus 488 (1:100, ThermoFisher, #A32931). Sections were mounted with Vectashield with DAPI (Vector Laboratories).

### Imaging and Analysis

Images were obtained in confocal microscope (Zeiss LSM 880) using a 20 X objective (NA:0.3) and all image quantification was performed using Fiji image processing package. Briefly, after splitting images in single channels, channels containing positive cells (eGFP, Cre or tomato positive) were threshold adjusted using the Huang algorithm and watershed transformation. Eight-bit transformed images were used as masks to create ROIs of positive cells. ROIs were overlapped to the c-Fos channel and c-Fos signals were quantified according to the mean gray intensity. c-Fos positive cells were defined as cells with mean gray value above 4 standard deviations from background signal, which was defined as the average of 20 random selected ROIs. Images of dorsomedial striatum of both hemispheres were taken (2-4 sections/animal, at least 4 animals/genotype). All images were taken within approximately similar sections across samples. For the Cre/GFP experiment, slices with less than 20 infected cells/field were excluded from the analysis. Analyses were performed by investigator blinded to the experimental groups.

### Electrophysiology

For excitability experiments, male and female D1-Lrrk2-KO;Drd1-tdTomato+/- and littermate control mice (8-12 weeks-old) were anesthetized with isoflurane and perfused transcardially with warm (33 °C) artificial cerebrospinal fluid (aCSF) containing the following (in mM): 124 NaCl, 2.5 KCl, 1.3 MgCl_2_, 2.5 CaCl_2_, 1.0 NaH_2_PO_4_, 26.2 NaHCO_3_, 20 D-glucose, 0.4 ascorbate and 3 kynurenic Acid. Brains were removed and placed in a vibratome (Leica). Sagittal brain slices (230 μm) were prepared in warm aCSF. Slices were incubated in warm (33 °C) 95%O_2_/5%CO_2_ oxygenated aCSF containing kyrurenic acid (3 mM) for at least 30 min and moved to room temperature (22-24 °C) until used. Slices containing DMS were then transferred to the recording chamber that was constantly perfused with warm (33 °C) 95%O_2_/5%CO_2_ oxygenated aCSF at the rate of 1.5-2 ml/min. Drd1-tdTomato positive D1-MSN in the DMS were visualized with a 40x water-immersion objective on an upright fluorescent microscope (BX51WI, Olympus USA) equipped with gradient contrast infrared optics. Data was acquired using pClamp 10 software, sampled at 50 kHz and filtered at 1 kHz. Analysis was performed with AxoGraphX (Axograph Scientific). Whole-cell voltage clamp recordings were made from D1-MSN with patch pipettes (2.0-3.5 MΩ) filled with an internal solution containing (in mM): 120 K-methylsulfonate, 20 KCl, 2 MgCl_2_, 10 HEPES, 0.2 K-EGTA, 4 Na-ATP, and 0.4 Na-GTP, pH 7.35, 290 mOsm. Neurons were current clamped and adjusted to keep Vm at - 90 mV. Membrane responses to 1 s current injection between 0-700 pA (100 pA increment) were collected using an Axopatch-200B amplifier (Molecular Devices). Paired pulse optogenetic and electrical stimulation (20 Hz every 30 s) were used to trigger GABA-A IPSCs in GABA neurons in SNr. Physiological identification of GABA neurons was based on the rate of spontaneous action potential rate (>5 Hz) with spike widths <1.2 ms. Whole cell voltage clamp recordings were made with patch pipettes (2-3.5 MΩ) filled with an internal solution containing (in mM): 57.5 KCl, 57.5 K-methylsulfate, 20 NaCl, 1.5 MgCl_2_, 5 HEPES, 10 BAPTA, 2 ATP, 0.2 GTP, and 10 phosphocreatine, pH 7.35, 290 mOsM. To isolate GABA-A currents, excitatory synaptic blockers, NBQX (5 μM) and 3-((*R*)-2-carboxypiperazin-4-yl)-propyl-1-phosphonic acid (CPP 5 μM) were added to aCSF. All neurons were voltage clamped at -60 mV. Series resistance was monitored throughout the experiment (range; 3-15 MΩ). A fiber optic (200 μm/0.22 NA) connected to a blue LED (470 nm; Thorlabs) and bipolar electrical stimulator were placed near the recording neuron, and light stimulation (0.2-2 ms) was given to evoke GABA-A IPSCs. Whole-cell voltage clamp recordings from D1-MSNs were made to measure AMPAR/NMDAR ratio using patch pipette (2-3.5 MΩ) filled with internal solution containing (in mM): 120 CsMeSO_3_, 10 CsCl, 10 HEPES, 0.2 EGTA, 4 ATP, 0.4 GTP, and 10 phosphocreatine, pH 7.35 (290 mOsM) at 25°C. Evoked EPSCs were generated by electrical stimulation near the recording neurons with a glass monopolar microelectrode filled with ACSF. EPSCs were recorded in presence of GABA-A receptor blocker gabazine (5 μM). Evoked AMPAR current was recorded at -60 mV and measured the peak amplitude at 3–6 ms following the stimulation. Evoked NMDAR current was measured at 40 mV in the presence of NBQX (5 μM) and measured the peak amplitude around 10-15 ms following the stimulation.

### Stereotaxic virus injections

Mice were anesthetized with isoflurane and placed in the stereotax (Kopf Instruments). Adeno-associated viral (AAV) vectors for ChR2 expression (rAAV5-CaMKIIhChR2(H134R)-EYFP; 1.5×10^13^, Penn Vector Core), Cre expression (pAAV.CMV.HI.eGFP-Cre.WPRE.SV40) and eGFP expression (pENN.AAV.CB7.CI.eGFP.WPRE.rBG) were bilaterally injected into DMS (276 nl, from bregma: AP, +1.1; ML, ±1.2; DV, -3.0 mm for ChR2 expression and AP, +0.6, +1.1; ML, ±1.4, ±1.2; DV, -3.2 mm for Cre and eGFP expression) using a Nanoject II (Drummond Scientific).

### Alcohol-inducing locomotion

For all behavioral tests mice were habituated to the testing room for at least 30 min before starting the tests. Locomotor activity was measured during the animal’s light cycle in a clear polycarbonate chamber (20 cm H x 17 cm W x 28 cm D) equipped with infrared photobeam detectors (Columbus Instruments). The locomotor test consisted of baseline locomotor recordings of one-hour followed by an extra hour of recordings following intraperitoneal injections of saline or alcohol in the appropriate dose. Mice were first habituated to handling and injections for 3 consecutive sessions then injected with 1g/kg and 2g/kg of alcohol (20% v/v) on different days in a counterbalanced fashion. The dose of 3g/kg alcohol was administered on the last day to minimize tolerance. Data was analyzed as beam breaks per 5 minutes and normalized by the third day of saline injection.

### SKF81297-inducing locomotion

A similar procedure was performed as described above. However, locomotor activity was assessed using the IR actimeter system (Panlab). Briefly, after 3 days of habituation to saline injection, mice were injected with SKF81297 (Hello Bio, HB1858, 2 mg/kg). Locomotor activity was recorded for one hour before injection followed by another hour post injection.

### Open field and novel object exploration

Exploratory behavior was measured as distance travelled in a large open-field acrylic/PVC chamber (40×40×40 cm). Mice were allowed to explore for 30 min and then returned to homecage for 5 min, during which time a novel object was placed in the center of the open field arena. Animals were placed back into chamber and allowed to explore for an additional 15 min while video recording. The mouse position was tracked over time to estimate distance travelled using EthoVision XT (Noldus).

### Loss of righting reflex

Mice were injected with alcohol (3 g/kg, 20 mL/kg, *i*.*p*.) and placed supine in a v-shaped trough. The loss of righting-reflex was defined as the mouse inability to right itself three times within a period of 30 s. The time to lose and the time to regain the righting-reflex was recorded. The maximum time to regain the righting reflex was capped to 60 min. If mice failed to respond within that time frame, the session was terminated and the regain time was defined to 60 minutes. Blood samples were collected at the time of regain of righting reflex and BAC was measured using AM1 Alcohol Analyser (Analox).

### Alcohol sensitization

Behavioral sensitization to the stimulant effects of alcohol, was adapted from previously described procedure (*38*). Locomotor activity was recorded in a polycarbonate chamber (20 cm H x 17 cm W x 28 cm D) equipped with infrared photobeam detectors (Panlab). Mice were first habituated to saline injections for two daily consecutive sessions then injected with alcohol (2g/kg, 12.5ml/kg) for 8 consecutive sessions/day. Locomotor activity was recorded for 15 minutes. Following the last day of alcohol injections, mice were left undisturbed in their home cage for 8 days followed by a single alcohol challenge session.

### Alcohol drinking behavior

#### Intermittent access two-bottle choice

mice had intermittent access (Mon-Wed-Fri) to one bottle with unsweet 20% alcohol solution in tap water and continuous access to another bottle with tap water in the home-cage for 4 weeks. The position of water and alcohol bottles were switched every drinking day to avoid side preference. Bottles were weighted before and after each 24h-session and volume consumed was calculated from the weight difference. Alcohol intake was calculated in grams of alcohol consumed per kilogram of body weight (g/kg). Alcohol preference was calculated as consumed volume of alcohol solution over the total consumed volume (water + alcohol solution).

#### Alcohol dose-response and quinine adulteration

this procedure was carried out as described for the 2-bottle choice paradigm, but animals had continuous access for 4 days/week to both bottles (Mon through Fri) for 7 weeks. Alcohol concentration was 20% during week 1 and 2, then changed to 10% during week 3, and changed to 5% alcohol during week 4. On week 5, mice returned to 20% alcohol access during 2 first days to regain baseline and then tested for their resistance to quinine adulteration by adding 0.25 mM quinine to the 20% alcohol solution for the last 2 days. On week 6, the same protocol was followed but 0.5 mM quinine was used instead. On week 7, water was adulterated with 0.25 mM quinine for 2 days and 0.5 mM quinine for 2 days for taste aversion testing. Alcohol consumption and alcohol preference were measured daily as described above.

#### Drinking in the dark (DID)

a modified drinking in the dark procedure was used according to previously published (*79*). Briefly, 3 h into the dark cycle (9:30 am) the water bottle in each cage was replaced by a new bottle containing 20% alcohol solution made available for 4 h. Procedure was repeated 5 days/week for 3 weeks. Alcohol intake was measured using change in bottle weight as explained above. Following DID procedure, mice were trained to self-administer alcohol in operant chambers.

#### Operant alcohol self-administration

*Acquisition of operant behavior:* the SIPPER training procedure was adapted from previously published study (*79*). Mice were trained in an operant chamber (Med Associates) to press a lever to gain access to a sipper tube that delivers a 20% alcohol solution. Each chamber contained two levers (active and inactive), a green cue light located on top of the active lever and a retractable sipper that was covered by an automatic door. The retractable sipper is attached to a 5 ml glass pipette containing the alcohol solution, which allowed for precise measurements of the consumed volume during the session. A lickometer attached to the sipper records contacts with a 10 ms time resolution. Chambers are housed in a sound-attenuating box coupled with a white-noise ventilation fan. Training sessions were 6h long and performed on intermittent days (2-3 sessions a week, no sessions on weekends). Mice were first trained at a fixed ratio 1 (FR1) for 9 sessions in which each press on the active lever resulted in sipper extension into the chamber for 60 seconds. Presses on the inactive lever or on the active lever during an ongoing access time were recorded but had no consequence. The cue light located above the active lever was illuminated as default and turned off after a successful ratio was achieved and remained on during sipper extension time. After the initial acquisition phase, the fixed ratio was increased to 3 (FR3) for 4 additional sessions. All mice showed less than 30% variation on the daily active response or alcohol intake by the last FR1 session. A food pellet was made available on the cage floor during all sessions. Mice that did not reach the minimum criteria of average alcohol intake of more than 0.1 g/kg alcohol were classified as non-drinkers and excluded from further analysis. *Progressive ratio session:* mice were tested in a single progressive ratio session in which the number of lever presses required to earn access to the sipper was increased exponentially (*80*). In addition to successfully completing the appropriated lever press, the progress to the next ratio was also contingent to a minimum of 10 lick contacts to the lixit sipper. Cases in which a successful lever press ratio was completed but the minimum lick requirement was insufficient, the current ratio was kept the same. Session was terminated after 1h of unsuccessful ratio progression or after a total session time of 5 h.

*Quinine adulteration session:* after the progressive ratio session, mice returned to the regular training session (FR3, 60s access time) until restauration of baseline levels of responding. After retraining, the alcohol solution was adulterated with 0.25mM quinine while the other parameters were kept the same. Testing was performed for two sessions and the impact of quinine adulteration on alcohol drinking was evaluated as the percentage of change in alcohol consumption during the quinine sessions compared to retraining sessions (alcohol consumption during quinine session / alcohol consumption during retraining session * 100).

*Foot shock sessions:* after quinine adulteration sessions, mice returned to regular training session (FR3, 60s access time) to recover baseline levels of responding. Three shock sessions were carried out as training sessions, except that every other alcohol access was paired with delivery of a foot-shock that lasted 0.5 seconds. Shock intensity was increased with each consecutive session from 0.2, 0.4 to 0.6 mA. Resistance to punishment was measured as the percentage change in alcohol consumption during the punished sessions compared to retraining sessions (alcohol consumption during foot shock sessions / alcohol consumption during retraining session * 100).

### Sucrose and sucralose preference test

Sucrose preference was measured using a two-bottle choice paradigm for 4 days. Naïve mice had continuous ad libitum access to water and intermittent access to sucrose for 6 hours/day. During days 1 and 2, mice had access to 1% sucrose in tap water and for day 3 and 4 to 2% sucrose in tap water. A second cohort of mice had also access to 0.5% sucrose after the dose of 2%. Sucralose preference was assessed in a similar fashion of the sucrose test. Sucralose concentrations were 1%, 0.5% and 0.05%, respectively. Preference was assessed as the percentage of sucrose solution consumed over the total fluid intake per 6 hours (solution consumption / solution consumption + water consumption * 100).

### Food consumption

Food consumption was recorded using two diets, regular chow (NIH 31, 4.7 kcal% fat; 3.0 Kcal/g) and high fat diet (HFD; D12492, 60 kcal% fat; 5.2 kcal/g; Research Diets, Inc). During experimental procedure, mice had 24h/day ad libitum access to regular chow for one week followed by HFD access for 2 weeks. Food was weighed every day at 10:00am and mice were weighed on the seventh day of each week. Food consumption was calculated as caloric intake multiplying the amount of food consumed by the caloric content of each diet.

### Hot plate test

During the habituation phase, mice (n=6/8) were allowed to freely explore the apparatus for 10 min. Mice were returned to the homecage for up to 2h and were placed back on the metal surface maintained at a constant temperature of 52.2 °C. The time taken to elicit licking of the forepaws was recorded as the latency and used to score pain sensitivity. Each mouse was tested three times with a minimum intertrial interval of 10 min. Latency in each trial were averaged to calculate individual scores.

### Drugs and chemicals

Alcohol (Decon Laboratories, 190 proof, glass container) was dissolved in tap water at 20% v/v for drinking experiments. Quinine hemisulfate monohydrate (Sigma-Aldrich) was dissolved in the 20% alcohol solution or in tap water. SKF81297, NBQX, and CPP were purchased from Tocris. SCH23390, gabazine, and kynurenic acid (sodium salt) were obtained from Abcam.

### Statistical Analysis and Quantification

The numbers of biological replicates, observations made, and the exact statistical tests used can be found in the relevant figure legends and in the text. Graphs and analyses were performed in Prism 7 (GraphPad), Igor Pro 9 (Wavemetrics) and R software (version 4.0.3). Unless stated otherwise, comparisons among two groups were performed using 2-tailed t-test or the non-parametric correspondent Mann-Whitney test, for paired or unpaired samples as appropriate. Three or more group comparisons were always analyzed using 2-tailed ANOVA, unless mentioned. When appropriate 2-way repeated measures ANOVA, Mixed-effects model ANOVA (REML) or 1-way ANOVA, were used. Unless otherwise stated, Sidak’s and Tukey’s test were used for post hoc multiple comparisons. Violation of sphericity in RM-ANOVA was corrected using the Greenhouse-Geisser method. Experimental design determined the statistical test used and all data met assumptions for the test. Results were considered significant at an alpha of 0.05. Data are presented as mean ± SEM. Detailed statistics are described in supplementary Table S1.

## Acknowledgments

We thank Dr. Huaibin Cai (NIA) for generously providing the Lrrk2 conditional mouse line and the members of the Alvarez laboratory for their valuable comments on the manuscript. The authors declare no competing financial interests. This work utilized the computational resources of the NIH HPC Biowulf cluster (http://hpc.nih.gov). We thank the NIH Fellows Editorial Board for proofreading the manuscript.

## Funding

ZIA-AA000421, Intramural Research Programs of National Institute on Alcohol Abuse and Alcoholism, NIH (to V.A.A)

UO1 AA023489, National Institute on Alcohol Abuse and Alcoholism, NIH (to D.R. and V.A.A)

National Institute on Aging, NIH (to MRC)

NIH Director Innovation Award, Center on Compulsive Behaviors, NIH (to VAA)

## Author contributions

Conceptualization: DDSES and VAA.

Research design and Methodology: DDSES and VAA

Behavioral, molecular and in silico analysis: DDSES

Electrophysiology experiments and analysis: A Matsui

Biochemistry experiments and analysis: A Mamais

RNAscope and immunochemistry experiments: DDSES and EM

Supervision of experiments and analysis: VAA, MRC, DR

Writing – original draft: DDSES and VAA

## Data and materials availability

The data that support the findings of this study are available from the corresponding author upon reasonable request.

**Fig S1:**
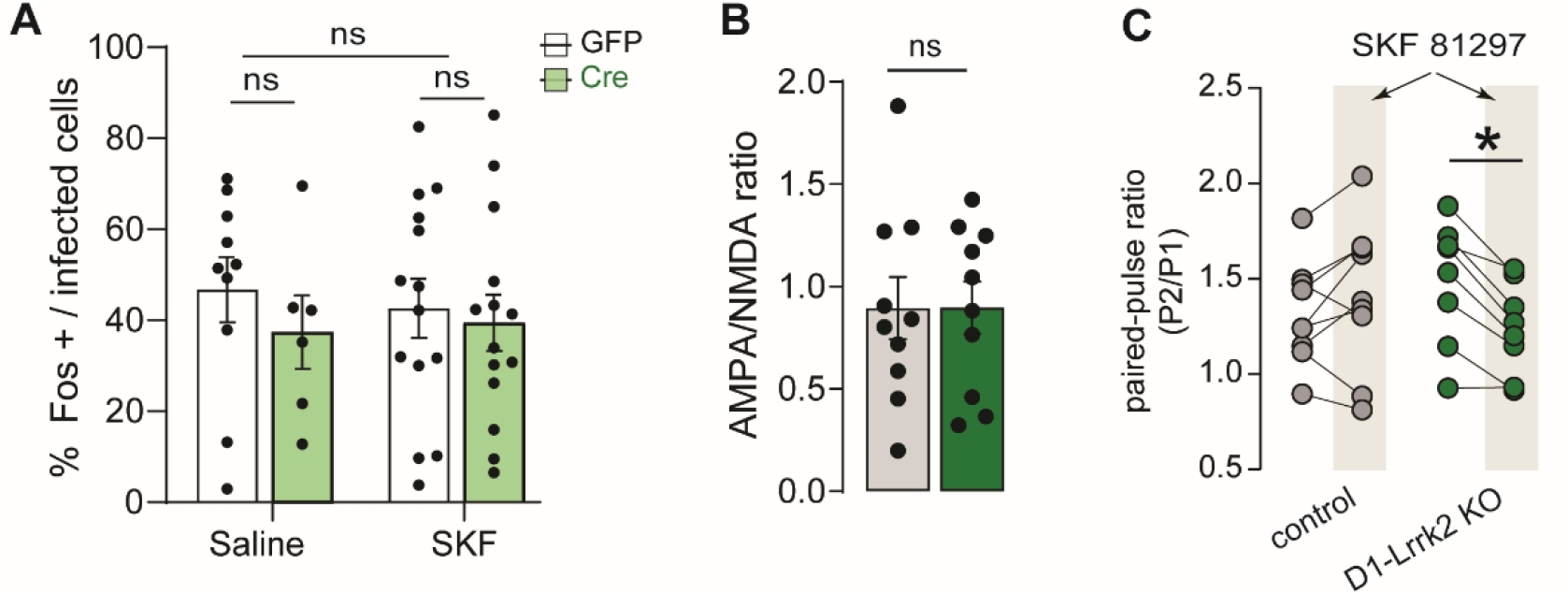
Assessment of D1R, Glutamatergic, and GABAergic function in D1-Lrrk2-KO mice. **A**, Quantification of c-Fos positive cells in the striatum of Lrrk2^floxP/floxP^ mice injected with either Cre(eGFP) or GFP after systemic administration of saline or SKF81297 (2 mg/kg). **B**, Average AMPAR/NMDAR ratio. **C**, Paired pulse ratio of the synaptic responses before and after (shaded area) bath application of D1-like agonist. For all panels, bars represent mean ± S.E.M and symbols represent values from individual mice. * denotes *P* < 0.05;

**Fig S2:**
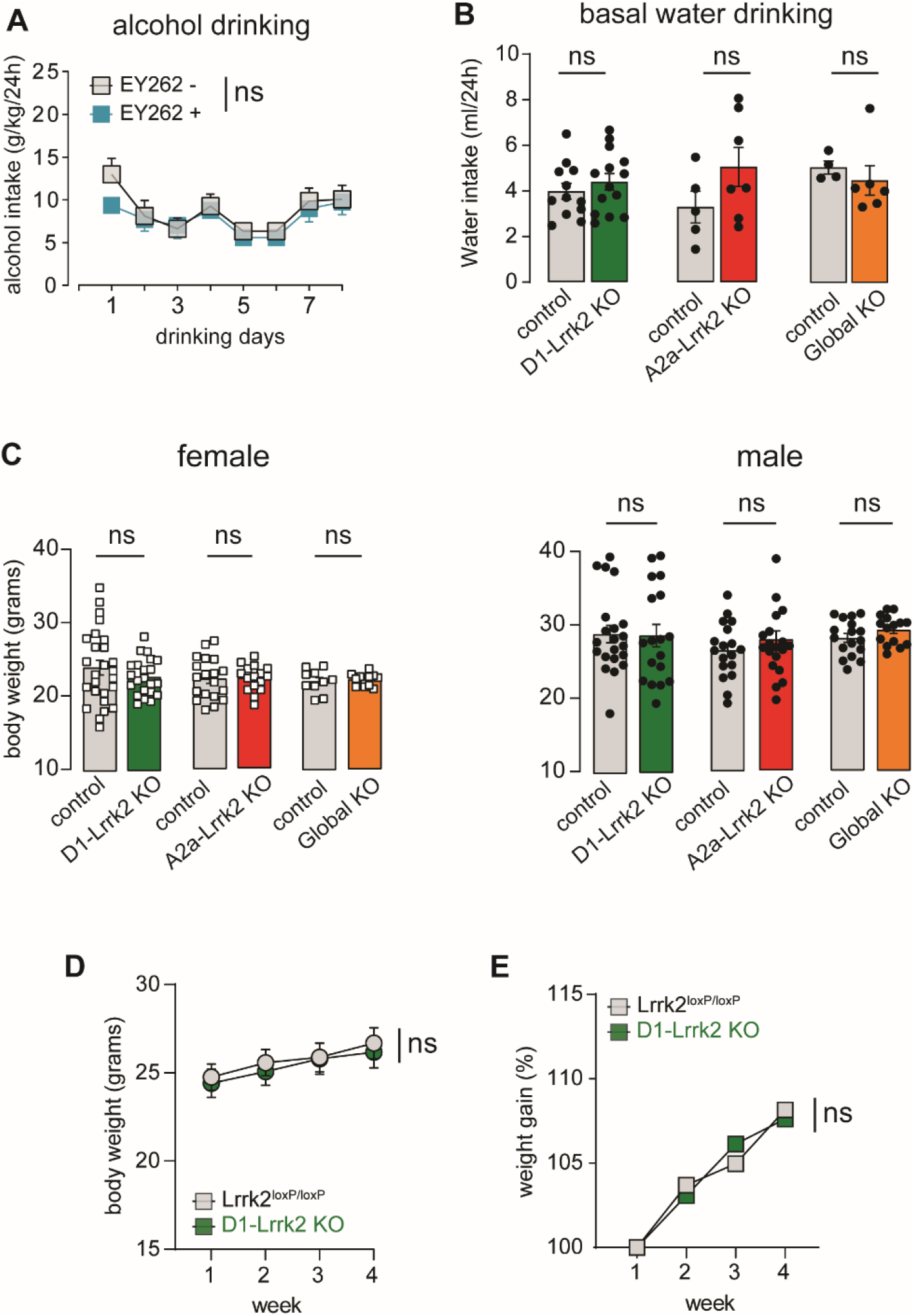
Additional data on consummatory behaviors in cell-specific Lrrk2-KO mice. **A**, Mean alcohol intake (g/kg/24h) during 2-bottle-choice sessions for EY262 (blue) and littermate controls (gray). **B**, Bar graphs showing basal water drinking (ml/24h) for D1-Lrrk2-KO (green), A2a-Lrrk2-KO (red), Global-Lrrk2-KO (orange) and their respective littermate controls (gray). **C**, Bar graphs showing average body weight for females (left) and males (right) for D1-Lrrk2-KO (green), A2a-Lrrk2-KO (red), Lrrk2-Global-KO (orange), and their respective littermate controls. **D-E**, Raw (D) and percentage (E) of weight gain for D1-Lrrk2-KO and controls during 4 weeks of drinking procedure. Bars represent mean ± S.E.M and symbols represent values from individual mice.

**Fig S3:**
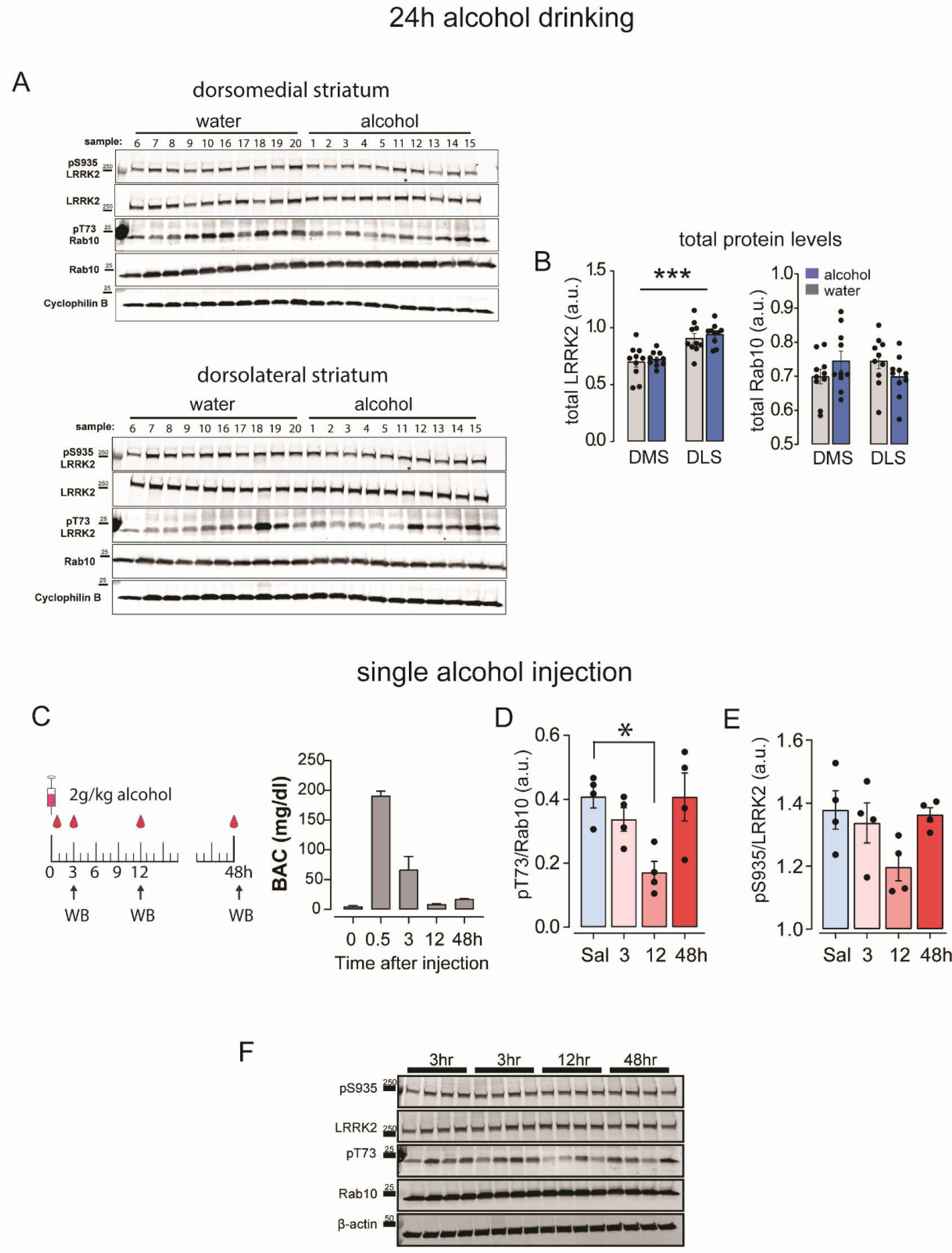
Additional data related to effects of alcohol on LRRK2 kinase activity. **A**, Western Blots for pS935 LRRK2, LRRK2, pT73 RAB10, RAB10, and Cyclophilin B in the DMS (*top*) and DLS (*bottom*) for alcohol and water drinking mice. **B**, Bar graphs showing labeling density for total LRRK2 (*left*) and total Rab10 (*right*) in the DMS and DLS of water (gray) and alcohol (blue) drinking mice. **C**, *Left*, Schematic diagram of systemic injections of 2g/kg alcohol and time points of tissue collections for Western blot. *Right*, Blood alcohol levels in different time points after systemic injection of 2g/kg alcohol. **D-E**, Time-dependent changes in pS935 LRRK2 (E) and pT73 RAB10 (F) levels after systemic injection of 2g/kg alcohol. **F**, Western Blots for pS935 LRRK2, LRRK2, pT73 RAB10, RAB10, and ß-actin in the DMS of mice injected with saline or 2g/kg alcohol. For all panels, bars represent mean ± S.E.M and symbols represent values from individual mice. * denotes *P* < 0.05, *** denotes *P* < 0.0001

**Fig S4:**
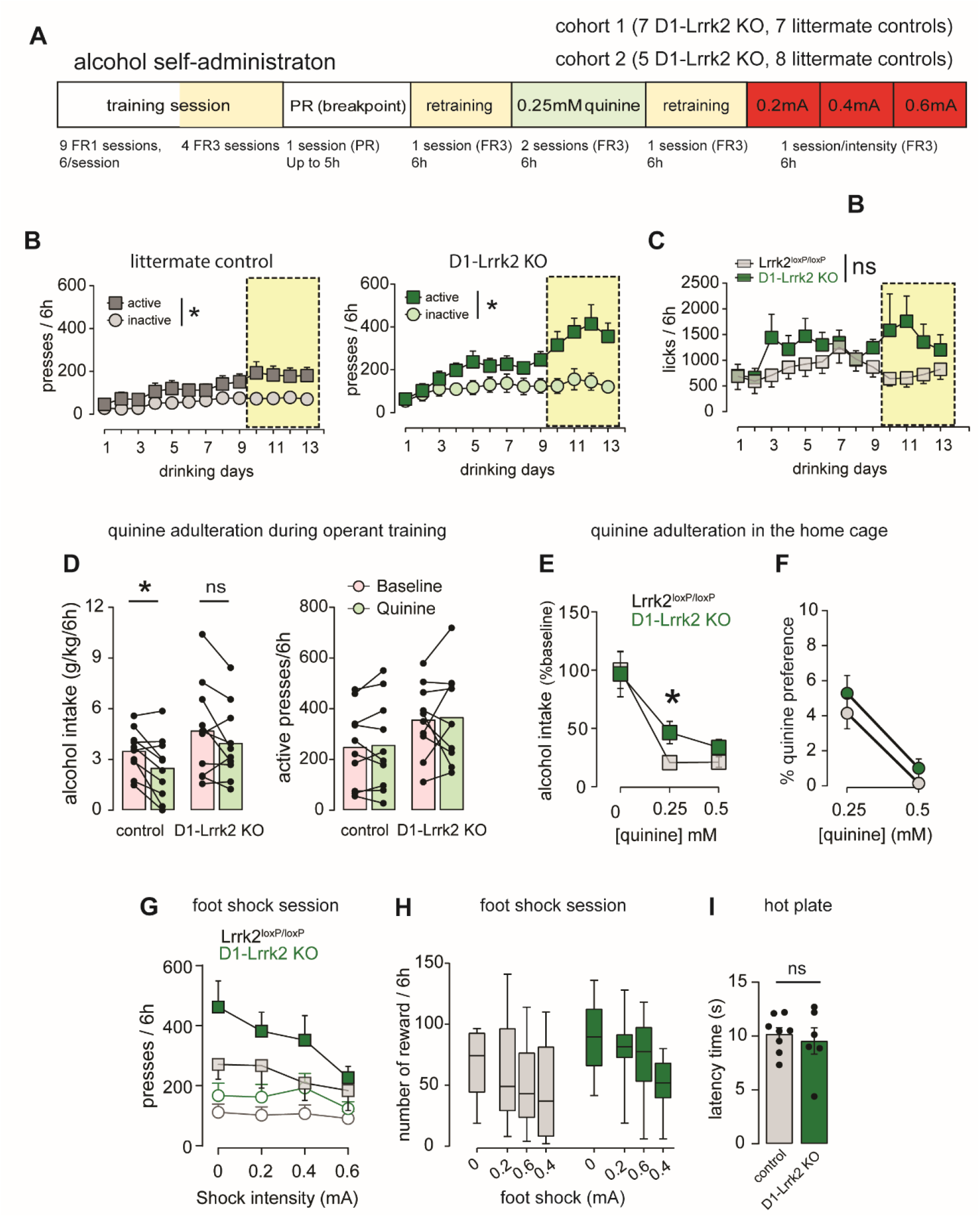
Operant alcohol self-administration in D1-Lrrk2-KO mice. **A**, Detailed schematic diagram of the operant training schedule. **B**, Mean number of active and inactive lever presses. **C**, Mean number of licks. **D**, Average alcohol intake (*left*) and active lever presses (right) during baseline (pink) and quinine adulteration (*green*) sessions. **E**, Non-operant alcohol intake (g/kg/24h) during baseline sessions with access to 20% alcohol solution (0) and during quinine-adulteration sessions. **F**, Quinine preference (taste aversion) tested using two bottle choice for tap water or quinine solution (0.25mM and 0.5 mM). **G**, Mean number of lever presses during foot shock sessions. **H**, Average number of earned rewards during baseline and foot shock sessions. **I**, Average withdrawal latency in the hot plate test. For all panels, bars represent mean ± S.E.M and symbols represent values from individual mice. * denotes *P* < 0.05

**Fig S5:**
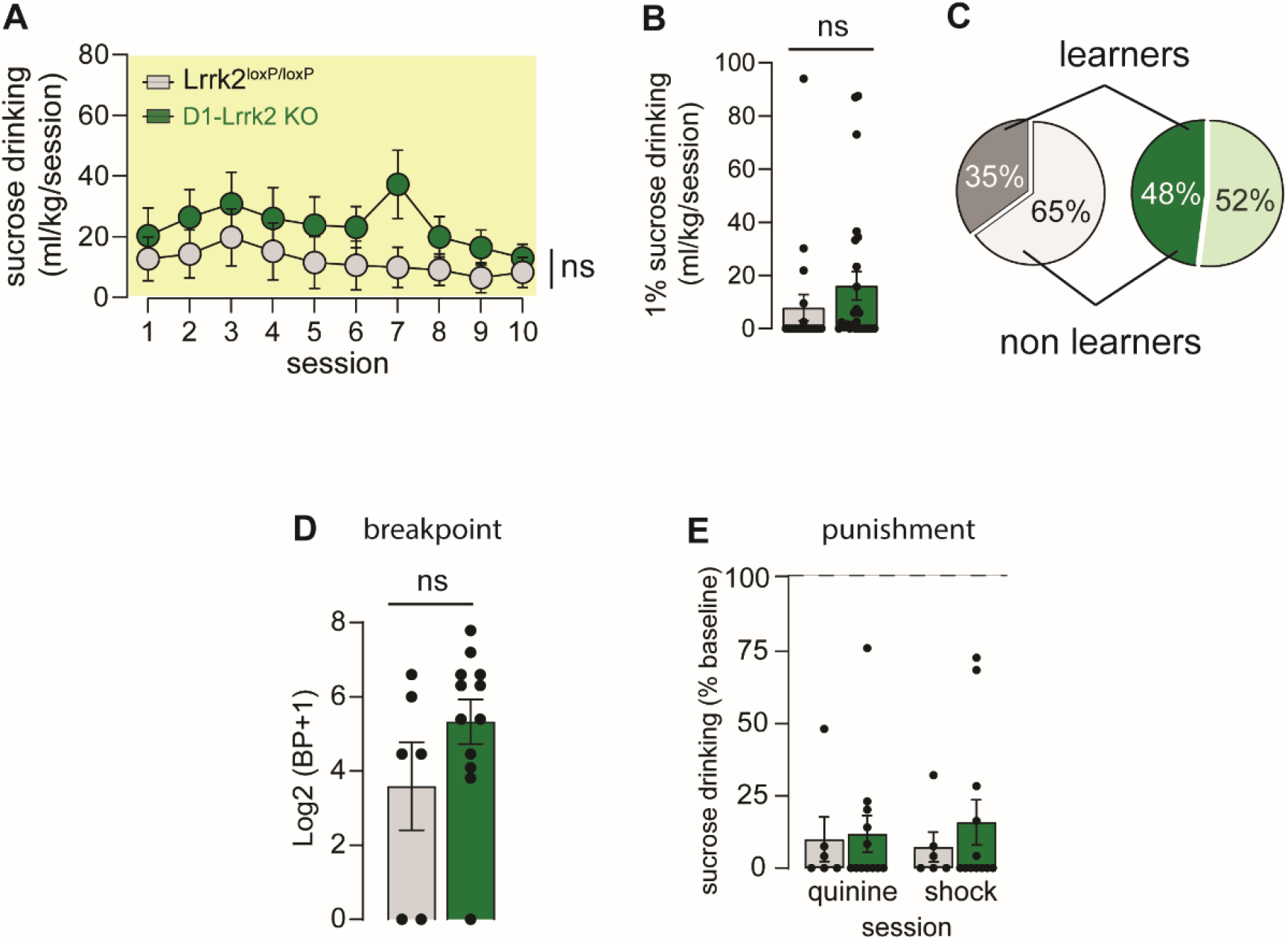
Operant sucrose self-administration in D1-Lrrk2-KO mice. **A**, Sucrose consumption during operant self-administration task. **B**, Average intake during access to 1% sucrose solution. **C**, Proportion of mice that acquired (learners) or did not acquire (non-learners) the self-administration task. **D**, Breakpoint for sucrose. **E**, Sucrose drinking during quinine adulteration and foot shock punishment sessions.

